# AGC kinase homology requires and enables co-targeting for CNS regeneration

**DOI:** 10.64898/2026.08.28.746753

**Authors:** Hassan Al-Ali, Abdiel Badillo-Martinez, Bassel Awada, Brian B. Silver, Omar S. Elwardany, Ronald N. Buckle, Graham Johnson, Siyuan Sun, Rodrigo Lendof, Nikhita Guhan, Praveen Singh, Jae K. Lee, John L. Bixby, Vance P. Lemmon

## Abstract

Axon regrowth in the central nervous system (CNS) is constrained by robust regulatory networks. Here we show that optimal neurite outgrowth in rodent and human CNS neurons is achieved by co-inhibition of kinases across four closely related clades within the protein kinase A, G, and C (AGC) family. The kinases derive from ancestral regulators of cytoskeletal dynamics, resource allocation, and polarized cell growth. Their shared domain architecture makes polypharmacology (co-engagement by a single small molecule) feasible. Phenotype-guided optimization of a tool compound with established efficacy in mouse spinal cord injury models yielded TMP-316, a drug candidate engaging these AGC kinases with selectivity against the broader kinome. A single intrathecal dose of TMP-316 produced sustained motor recovery in a rat cervical hemicontusion model. These findings reveal conditions under which polypharmacology is simultaneously required and enabled by shared evolutionary origins, illuminating a therapeutic discovery principle for pathologies governed by functionally overlapping targets.

---

Spinal cord injury (SCI) causes permanent loss of motor and sensory functions, and decades of research have failed to produce an approved therapy to promote regeneration and functional recovery^1^. This failure reflects a convergence of multiple repressive mechanisms that restrict the regenerative capacity of injured central nervous system (CNS) neurons. Genetic studies have demonstrated that the regenerative machinery remains latent and can be reactivated^2–4^, but this insight has yet to be translated into a clinical strategy.

Therapeutic development in SCI has overwhelmingly followed the single-target paradigm^5^. Disorders featuring biological robustness, where compensatory coverage among overlapping regulatory networks buffers single-node perturbation, have consistently resisted single-target intervention^6,7^. Multi-target agents now account for a substantial fraction of approved therapeutics^8^, yet no theoretical framework determines when multi-target engagement is required or which targets must be coordinated. Our prior studies, combining phenotypic and target-based screening with machine learning^9–11^, identified multiple members of the protein kinase A, G, and C (AGC) family as potential targets for promoting axon regeneration^9^. Several of these kinases were previously pursued as single targets^12,13^, but their functional coordination was unrecognized, and the polypharmacology of common kinase inhibitors likely led to misattributed efficacy in some cases.

Here, we demonstrate that optimal neurite outgrowth requires concurrent inhibition of kinases from four related AGC clusters. Mechanistic studies using S6K1, ROCK2, PKCγ, and PKX as cluster representatives establish their additive contributions to neurite outgrowth. Evolutionary analysis traces this coordinated action to conserved ancestral functions and structural analysis shows that homologous active-site architecture makes single-agent engagement structurally feasible. RO48, a tool compound with polypharmacology across these kinases, previously showed histological and functional efficacy in mouse SCI models ^9,14^. Phenotype-guided structure-activity-relationship (SAR) studies of RO48 yielded TMP-316, a drug candidate that retains AGC kinase polypharmacology while exceeding the kinome-wide selectivity of most FDA-approved kinase inhibitors. A single intrathecal (IT) dose of TMP-316 produced sustained motor recovery through the six-week study endpoint in a rat cervical hemicontusion model.

## Phenotype-guided SAR produces TMP-316

We previously developed idTRAX, a target deconvolution platform combining phenotypic screening with biochemical profiling and machine learning^9,10^. It identified several AGC kinase groups as potential repressors of neurite outgrowth (Extended Data Table 1), with siRNA knockdown providing in vitro validation in primary neurons^9^. RO48, a kinase inhibitor with polypharmacology spanning these groups, promoted the greatest neurite outgrowth of all compounds in our prior campaigns^9,15^. A single intracortical injection of RO48 promoted regeneration and functional recovery in mouse models of SCI^14^. RO48 exhibits hERG activity and a short CSF half-life, limiting translational utility^9,14^. We therefore sought to develop a therapeutic candidate deliverable via a clinically feasible route. Lumbar IT delivery was selected for two reasons: lumbar catheterization is routinely performed in SCI patients to manage cerebrospinal fluid pressure^16^, and IT administration directs drug to the CNS while minimizing systemic exposure^17^.

To avoid constraining chemistry to an incomplete list of targets, a hit-to-lead campaign was guided by neurite outgrowth SAR rather than optimization against individual targets. Compounds were rank-ordered using the percent drug effect score (%DES)^18^, a composite metric integrating potency and effect size, benchmarked to RO48 in each run. This captures each compound’s cellular activity while minimizing inter-run variability, which is necessary for reliably driving SAR with gain-of-function assays such as neurite outgrowth^18^. Activity against ROCK2 and S6K1 was tracked in parallel using in-cell engagement assays^19^ to ensure retention of these validated targets^20,21^. Medicinal chemistry pursued three axes: synthetic tractability, Pgp efflux reduction, and elimination of hERG activity and other liabilities (Methods). The effort spanned 371 analogs and yielded TMP-316 (BPN-0037316) (Fig. 1a, b; Extended Data Table 2).

**Figure 1.**
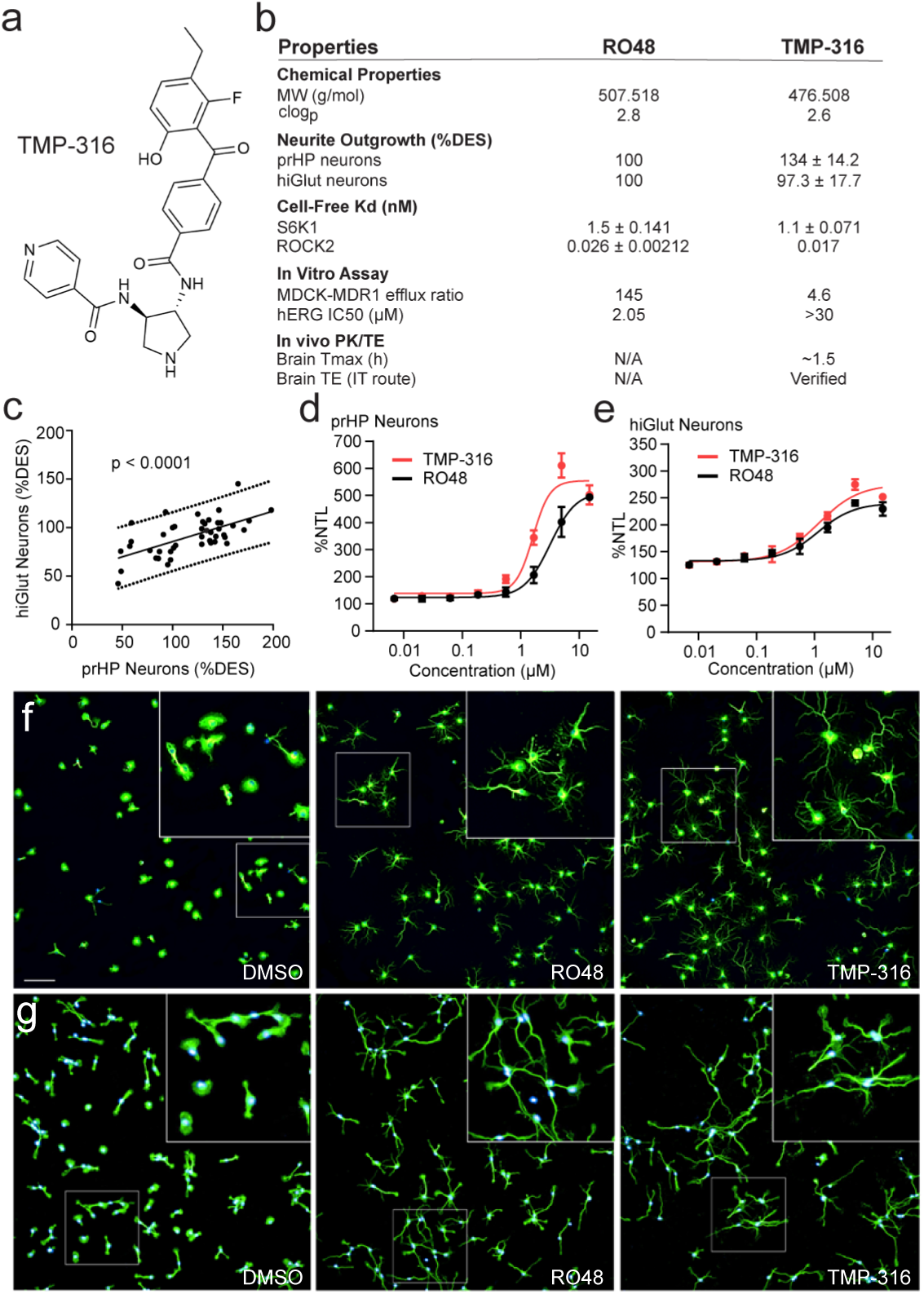
| TMP-316 is an analog of RO48 with improved properties. (**a**) Molecular structure of TMP-316. (**b**) Chemical and biological properties of RO48 and TMP-316. Values are presented as mean ± SD. (**c**) Correlation between %DES of BPN program compounds tested in primary rat hippocampal (prHP) and human iPSC-derived glutamatergic (hiGlut) neurons (N = 44, R2=0.4). **(d)** Neurite outgrowth (%NTL) of prHP and **(e)** hiGlut neurons treated with RO48 or TMP-316. Data are presented as mean ± SD (n = 3-6 technical replicates). Representative fluorescence images of (**f**) prHP and (**g**) hiGlut neurons treated with DMSO vehicle, RO48 [5 µM], or TMP-316 [5 µM] for 48 h. Blue: Hoechst (nuclei), Green: βIII-tubulin (soma and neurites). Scale bar: 100 µm.

We previously showed that neurite outgrowth responses to pharmacological treatments can vary across neuronal cell types^18^, with compounds active in some neuronal types and inactive in others, reflecting differences in mode of action (MoA) and neuronal biology. Establishing MoA conservation between the rodent model and human CNS neurons for this chemotype was therefore essential. Program compounds that were tested in both primary rat hippocampal (prHP) and human iPSC-derived glutamatergic (hiGlut) neurons showed significantly correlated activities across the two cell types (Fig. 1c), consistent with verified neuronal expression and sequence similarity of AGC kinases in rodents and humans (Extended Data Table 1). The wider dynamic range and greater sensitivity of the rodent assay (Supplementary Fig. 1a, b), reflected in the regression slope skewing toward the rodent axis (Fig. 1c), made it the platform of choice for SAR, and the human assay was used to verify activity of program compounds and ensure translatability. Across both models, TMP-316 robustly promoted neurite outgrowth (Fig. 1d-g).

Reduction of the MDCK-MDR1 efflux ratio from 145 in RO48 to 4.6 in TMP-316 (Fig. 1b) was accompanied by substantial exposure in CSF, spinal cord, and brain following lumbar IT delivery in rats (Extended Data Table 3). A single IT dose of TMP-316 (∼1 mg/kg) detectably reduced cortical S6 phosphorylation at S240/244 (Supplementary Fig. 1c). This dose did not alter GFAP immunoreactivity or Iba-1+ cell density in the spinal cord (Supplementary Fig. 2), suggesting no adverse inflammatory effects even at the site experiencing the highest concentration of compound. TMP-316 lacks RO48’s potent hERG inhibition (Fig. 1b), addressing a key cardiac liability. Kinome-wide profiling of TMP-316 revealed that its activity concentrates within the AGC Kinase family (Fig. 2a), avoids several non-AGC off-targets engaged by RO48 (Supplementary Fig 3), and exhibits a Gini coefficient of selectivity of ∼0.78, exceeding that of many FDA-approved kinase inhibitors^22^.

**Figure 2.**
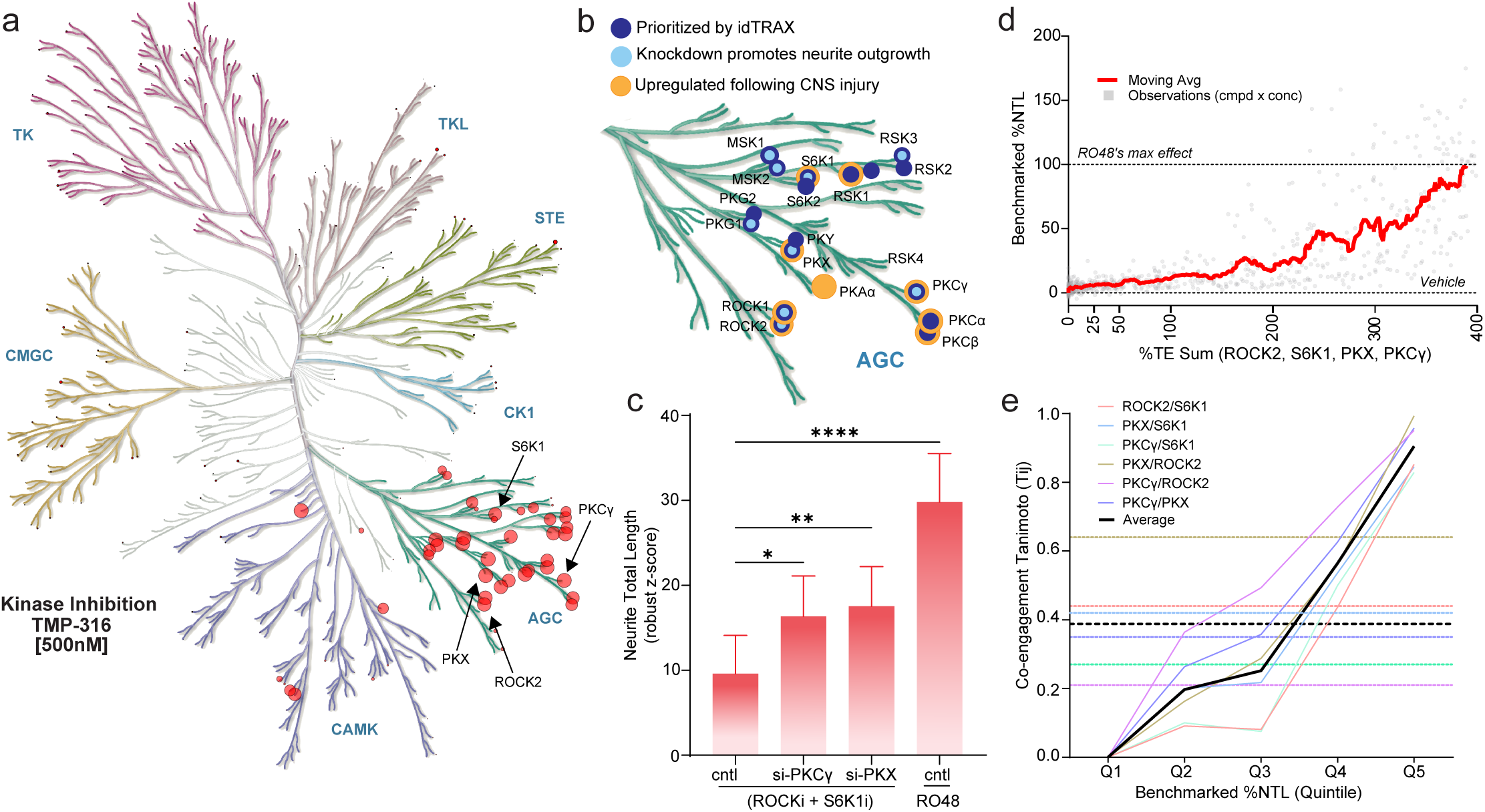
| Co-engagement of ROCK2, S6K1, PKCγ, and PKX drives neurite outgrowth. (**a**) Kinome selectivity profile of TMP-316. Node size scales proportionately with percent inhibition, with the largest node corresponding to complete inhibition. Illustration reproduced courtesy of Cell Signaling Technology, Inc. (www.cellsignal.com). Full inhibition data is provided in Supplementary Table 1 and the annotated kinome tree is available in Supplementary Figure 3b. (**b**) AGC kinases prioritized by idTRAX as targets for promoting neurite outgrowth, shown to promote neurite outgrowth in primary CNS neurons when silenced, or found to be upregulated or activated by CNS injury (Full data in Extended Data Tables 1 and 4). (**c**) Neurite outgrowth in prHP neurons transfected with scramble (cntl), α-PKCγ, or α-PKX siRNA and treated with ROCKi [10 µM] + S6K1i [5 µM] or RO48 [2 µM]. Data are presented as mean ± SEM (N = 4 biological replicates; one-way ANOVA with Dunnett’s multiple comparison test, * *p* < 0.05, ** *p* < 0.01, **** *p* < 0.0001). (**d**) Moving average of benchmarked %NTL values (prHP neurons) plotted against %TE values; n = 711 total observations each relating benchmarked %NTL to the sum of %TE S6K1, ROCK2, PKX, and PKCγ. (**e**) Co-engagement Tanimoto coefficient (Tij) for all six kinase pairs across Benchmarked %NTL quintiles (Q1–Q5; n = 142–143 observations per quintile, pooled across all compounds and concentrations). Solid colored lines represent per-pair Tij values calculated within each quintile; the solid black line represents the cross-pair mean. Dashed horizontal lines indicate the corresponding kinome-wide background Tij for each pair (colored) and for the cross-pair mean (black).

## Target co-engagement drives efficacy

Pharmacological linkage among closely related kinases precludes single-target deconvolution from small-molecule phenotypic screens; instead, idTRAX resolves targets as pharmacologically linked groups^9^. IdTRAX identified several AGC kinase groups as repressors of neurite outgrowth: S6K1/2, RSK1-4, MSK1/2, ROCK1/2, PKCα/β/γ, PKX, and PKG1/2 (Extended Data Table 1). These groups belong to four kinase clades^23^, a partition we adopt to operationally arrange them into four clusters (Fig. 2b, Extended Data Table 1). Several of these kinases are upregulated or activated by CNS injury (Fig. 2b, Extended Data Table 4), implicating them in the injury response. To tractably interrogate the causal relationship between co-inhibition and neurite outgrowth, one representative per cluster was selected for combinatorial studies. S6K1, ROCK2, PKCγ, and PKX were selected as representatives on the basis of convergent evidence: each was identified by idTRAX, its knockdown promoted neurite outgrowth, and prior studies indicate its upregulation or activation following CNS injury (Fig. 2b).

Selective small molecule inhibitors are not available for PKCγ or PKX, and combinatorial siRNA knockdowns of two or more targets often cause cytotoxicity that confounds neurite outgrowth assays. Therefore, siRNA was used to sequentially test whether knockdown of either PKCγ or PKX (Accell siRNA pools; knockdown efficiencies in Supplementary Fig. 1d, e) promotes neurite outgrowth beyond saturating co-inhibition of S6K1 and ROCK2. Bliss synergy analysis was used to identify the combined concentrations at which S6K1i and ROCKi saturate neurite outgrowth (Supplementary Fig. 1f, g). siRNA-mediated knockdown of either PKCγ or PKX promoted neurite outgrowth beyond that of the maximally active S6K1i-ROCKi combination (Fig. 2c).

The chemistry program produced 371 analogs with a range of activities on neurite outgrowth and differential selectivity across S6K1, PKX, ROCK2, and PKCγ (Extended Data Table 6). The availability of in-cell target engagement data for more than 75 of these created an opportunity to model the relationship between target engagement and cellular effect (Supplementary Fig. 1h, i). S6K1 inhibitor PF-4708671 ^24^ (hereafter S6K1i) and Rho Kinase inhibitor ROCK iIV^25^ (hereafter ROCKi) were included to anchor the model with the available selective inhibitors. The moving average of neurite outgrowth tracked with target co-engagement, with RO48’s maximal effect coinciding with full engagement of all 4 targets (Fig. 2d). Pairwise Tanimoto co-engagement (*T_ij_*) across the four kinases scaled monotonically with neurite outgrowth promotion, substantially exceeding background where neurite outgrowth is highest (Fig. 2e). Random Forest modeling reflects this relationship: prediction accuracy for neurite outgrowth increases as each representative kinase is added (Supplementary Fig. 1j, Extended Data Table 5). These results demonstrate that co-targeting these kinases additively promotes neurite outgrowth, with maximal growth directly correlating with full engagement of all four.

## Shared ancestry enables polypharmacology

The four clusters share a long evolutionary history that precedes the emergence of neurons. In unicellular eukaryotes, ancestral homologs function in polarized cellular extension, cytoskeletal dynamics, membrane integrity, resource allocation, and size control^26–28^. Subsequent metazoan evolution is marked by expansion and diversification of these lineages (Fig. 3a). To assess functional overlap among the clusters, we compared substrate preference profiles across 52 AGC kinases using percentile scores derived from more than 89,000 short peptide substrates^29^. Substrate preference represents only one determinant of kinase function; cellular specificity is further shaped by localization signals, scaffolding interactions, and regulatory domains not captured by this analysis. Nevertheless, as the most broadly available and phylogenetically unbiased proxy for functional similarity, this approach provides a principled strategy for assessing substrate overlap among these kinases. Pearson correlation of these profiles revealed that most kinases in the four clusters form a coherent block with broadly similar substrate preferences, separating from GRKs and PDK1 (Fig. 3b, Supplementary Fig. 4). Within this block, partial sub-blocks corresponding to known phylogenetic groupings (S6Ks, ROCKs, PKCs, PKA/PKX) were apparent but remained highly intercorrelated.

**Figure 3.**
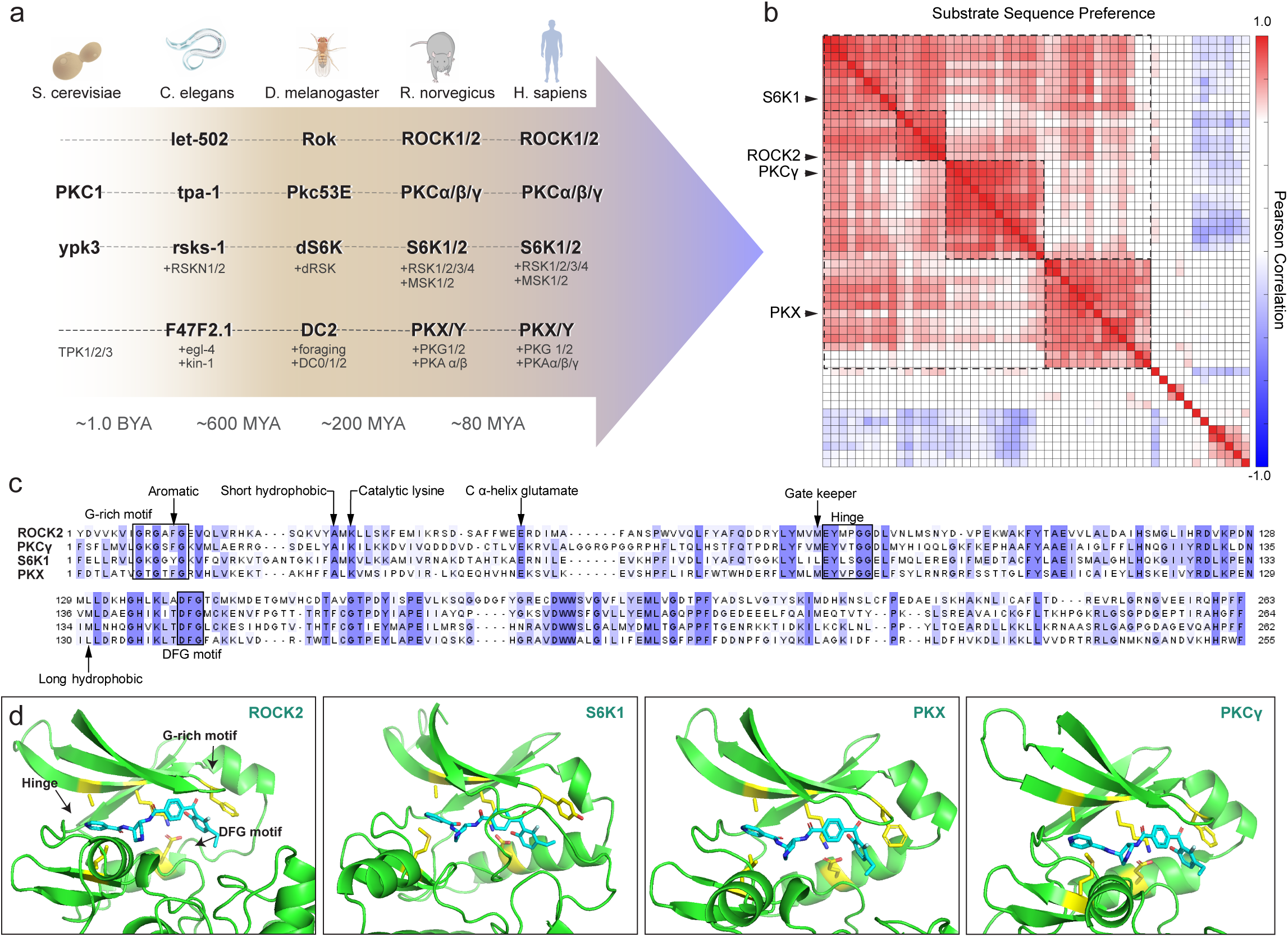
| AGC kinase homology underlies TMP-316’s polypharmacology. (**a**) Evolutionary lineage of AGC kinase clusters targeted by TMP-316, traced from unicellular eukaryotes to humans. Orthologs are shown across *Saccharomyces cerevisiae*, *Caenorhabditis elegans*, *Drosophila melanogaster*, *Rattus norvegicus*, and *Homo sapiens*. (**b**) Pairwise Pearson correlation of substrate sequence preference profiles across AGC kinases. Dashed boxes indicate blocks with similar substrate preferences. (**c**) Multiple sequence alignment of ROCK2, PKCγ, S6K1, and PKX kinase domains, with conserved structural motifs annotated (G-rich motif, DFG motif, hinge region, catalytic lysine, C α-helix glutamate, gatekeeper, aromatic, short hydrophobic, and long hydrophobic residues). (**d**) Co-folded structures of TMP-316 (cyan) within the kinase domains of ROCK2, S6K1, PKX, and PKCγ. Yellow sticks indicate conserved interacting residues within contact distance of TMP-316, comprising the catalytic lysine, C α-helix glutamate, aromatic residue of the G-rich loop, and the short and long hydrophobic residues flanking the hinge-binding region.

Shared ancestry also underlies conserved active site architecture, enabling single-molecule engagement. TMP-316 docks into the ATP-binding cleft of all four representatives (ROCK2, S6K1, PKX, PKCγ) with a similar type I binding pose engaging conserved hinge, G-rich, and DFG-in residues. Sequence alignment confirmed conservation of drug-interacting residues across all four (Fig. 3c, d). Conservation makes co-engagement structurally feasible but not inevitable: selective inhibitors and many program analogs achieve selectivity across these targets (Extended Data Table 6), indicating TMP-316’s polypharmacology reflects SAR-driven optimization, not passive consequence.

## Multitarget engagement remodels signaling

Quantitative phosphoproteomic profiling was performed in prHP neurons after treatment with TMP-316, RO48, S6K1i, ROCKi, or S6K1i+ROCKi, for 2 or 24 hours. Differentially phosphorylated proteins were mapped to Gene Ontology terms and sorted by functional category. Global pathway analysis identified biological processes overrepresented among treatment-responsive phosphoproteins, including categories related to microtubule cytoskeleton, growth cone and presynaptic function, actin cytoskeleton, neurite outgrowth, translation control, and autophagy (Fig. 4a, b, Extended Data Table 7, Supplementary Tables 4-8). PI3K/PTEN signaling showed stronger enrichment in RO48 and TMP-316 treatment.

**Figure 4.**
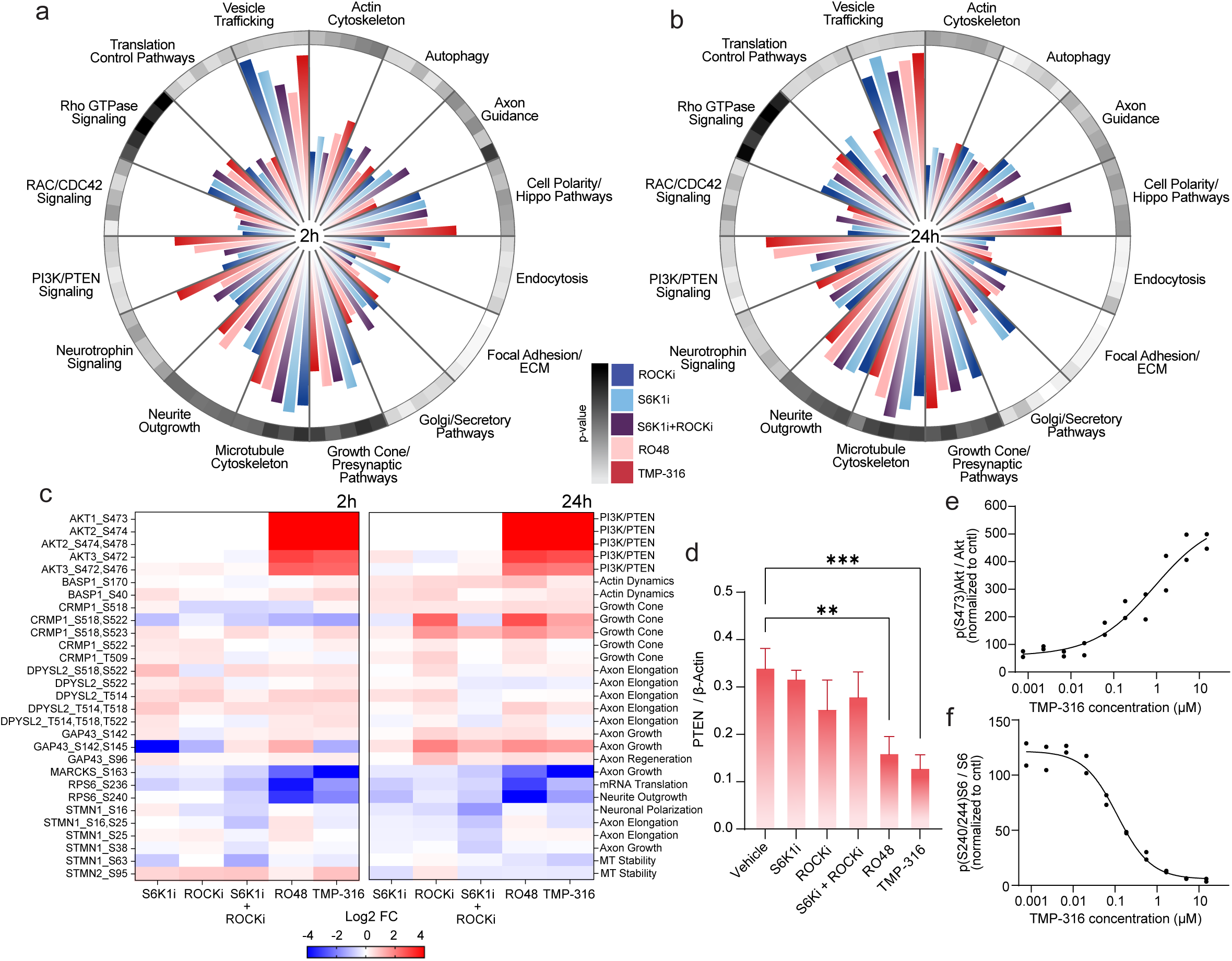
| RO48 and TMP-316 remodel neuronal signaling to drive neurite outgrowth. Radial bar charts depict pathway enrichment across 16 biological categories upon treatment with ROCKi, S6K1i, S6K1i + ROCKi, RO48, or TMP-316 relative to vehicle at **(a)** 2 h and **(b)** 24 h (n = 3 biological replicates). Bar length reflects the adjusted mean enrichment score per category. Ring heatmap depicts the mean p-value per category. **(c)** Heatmap of log2 FC relative to vehicle for selected phosphosites with established functional consequences that inform the directionality of selected pathways and biological processes identified in (a) and (b). Phosphosite details are summarized in Extended Data Table 8. **(d)** PTEN protein levels quantified by western blot from lysates of neurons treated with vehicle, S6K1i, ROCKi, S6K1i + ROCKi, RO48, or TMP-316. Data are presented as mean ± SD (N = 3 biological replicates; one-way ANOVA with Dunnett’s post-hoc test; ** p < 0.01, *** p < 0.001). Confirmation of TMP-316’s effects on **(e)** Akt (S473) and **(f)** S6 (S240/244) phosphorylation by western blot. Data are presented as mean ± SD; n = 2 technical replicates.

Because enrichment does not indicate pathway activation or inhibition, directionality was inferred separately from individual phosphosites with established functional consequence (Fig. 4c). RPS6 dephosphorylation at S236/S240 across all S6K1i conditions confirmed on-target S6K1 engagement. AKT phosphorylation reflected known S6K1 negative feedback on PI3K^30,31^ and ROCK-dependent PTEN regulation^32^. RO48 and TMP-316 produced stronger and longer-lived AKT hyperphosphorylation than the selective inhibitors or their combination. Western blot showed that TMP-316 and RO48 reduce canonical PTEN protein levels (Fig. 4d-f, Supplementary Fig. 5), suggesting an additional mechanism for activating PI3K signaling. PAK1/PAK2 phosphorylation reflected Rac1/Cdc42 activation^33^. MARCKS S163^34^, a canonical PKC substrate site, was selectively dephosphorylated by RO48 and TMP-316 but not by S6K1i, ROCKi, or their combination, marking a PKC-dependent signature absent from the dual-target combination. Additional regulated sites, including Stathmin, GAP43, and BASP1, and their functional significance are enumerated in Extended Data Table 8.

## Co-exposing soma and axons boosts growth

Phosphoproteomic profiling implicated mechanistic processes in both somatic (e.g., transcriptional regulation) and axonal (e.g., growth cone dynamics) compartments. Given that members of the target kinase clusters are expressed in both somatodendritic and axonal compartments of the neuron (Extended Data Table 1), we tested whether concurrent engagement of targets in both compartments is required for maximal outgrowth. RO48 was applied to the somatic compartment, the axonal compartment, or both, in prHP neurons cultured in microfluidic axon isolation chambers. Treatment of both compartments produced greater outgrowth than either somatic or axonal treatment alone (Fig. 5a, Supplementary Fig. 6), demonstrating that target engagement across both soma and axons leads to maximal outgrowth.

**Figure 5.**
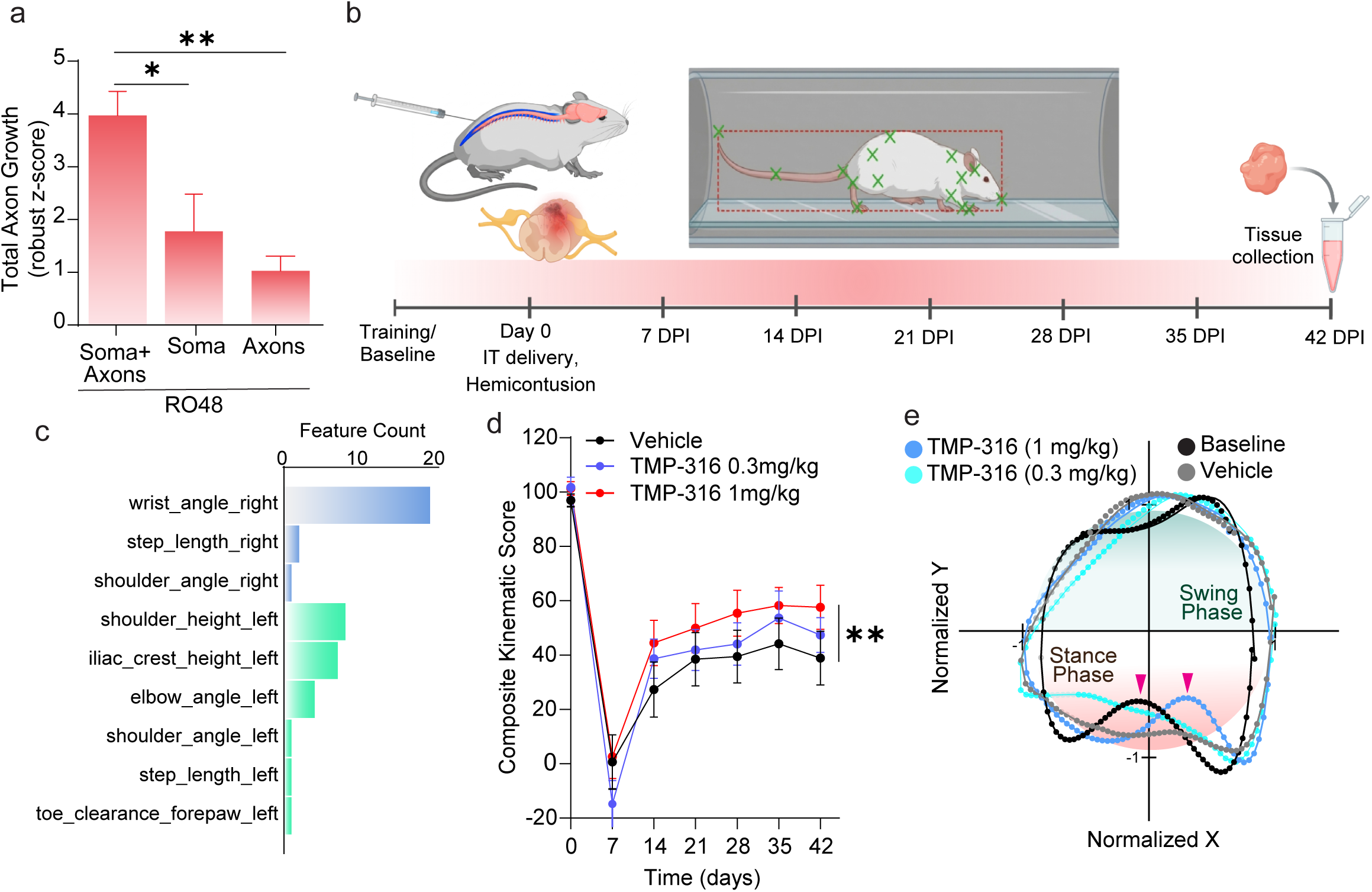
| Single intrathecal dose of TMP-316 promotes locomotor recovery in a rat cervical hemicontusion model. **(a)** rpHP neurons cultured in microfluidic chambers were axotomized at 8 DIV and treated with RO48 (2 µM) delivered to axons only, soma only, or soma and axons combined for 48 h. Data are presented as mean ± SEM (n = 4 biological replicates; one-way ANOVA with Tukey’s multiple comparisons test; * p < 0.05, ** p < 0.01). **(b)** Experimental timeline. Adult male Sprague Dawley rats were trained on the MotoRater device and baseline kinematics recorded prior to receiving a C5–C6 dorsal hemicontusion (170 kdyn) and a single IT dose of TMP-316 (0.3 or 1 mg/kg) or vehicle. MotoRater kinematic recordings were conducted weekly from 7 to 42 days post-injury (DPI); tissues were collected at 42 DPI. Created in BioRender. Silver, B. (2026) https://BioRender.com/he71yjw. (**c**) Number of selected kinematic features per base parameter (Methods), following two-stage TSFresh relevance filtering and ANOVA-based recovery-slope selection. (**d**) Composite Kinematic Score (CKS) over time for vehicle, TMP-316 0.3 mg/kg, and TMP-316 1 mg/kg groups. Data are presented as mean ± SEM (N = 12–13 per group; One-Way Repeated Measures ANOVA with Dunnett’s multiple comparisons test; ** p < 0.01). (**e**) Mean group toe-to-ear traces at 42 DPI, circularly normalized with swing and stance phases annotated. Pink arrowheads indicate point of maximum weight-bearing, identified by minimum toe-to-ear distance during stance phase.

## One dose produces sustained recovery

Prior experiments with mice in two different labs demonstrated that single intraparenchymal injections of RO48 in sensorimotor cortex were sufficient to produce substantial axon growth and behavioral recovery after transection injuries^14^. We therefore asked whether TMP-316, administered IT, would produce functional recovery in rats after cervical hemicontusion^35^, a model that matches most clinical SCIs^36^. We produced C5–C6 right cervical hemicontusion injuries in rats with an Infinite Horizon impactor. A single IT lumbar bolus of TMP-316 (1 or 0.3 mg/kg) was administered near the time of injury, and animals were assessed weekly for locomotor recovery from 7 to 42 days post-injury (DPI) (Fig. 5b). Histological analysis confirmed no significant differences in lesion size between groups (Supplementary Fig. 7). Multi-angle gait was recorded using a MotoRater, which extracts kinematic parameters from markerless tracking. Unbiased feature selection prioritized kinematic variables with clear relevance to cervical SCI, including toe clearance, forelimb joint angles, and ipsilateral shoulder height, along with contralateral limb and trunk features consistent with compensatory adaptations (Supplementary Fig. 8a). Recovery was quantified using the Composite Kinematic Score (CKS), a metric derived from principal component analysis (Methods). Animals treated with 1 mg/kg TMP-316 showed greater recovery than vehicle controls, with improvement starting at 14 DPI and the largest effect at 28–42 DPI (Fig. 5d, e). The 0.3 mg/kg group showed a non-significant recovery trend.

To verify the robustness of these results, recovery was quantified using three additional mathematically distinct metrics. All methods independently detected the high-dose effect (Supplementary Fig. 8b-d) and produced concordant trajectories, with maximal separation between high-dose and vehicle groups at 28–42 DPI. Detailed kinematic analysis of the injured forelimb revealed the functional basis of this recovery. Animals treated with 1 mg/kg TMP-316 showed improved weight-bearing on the injured right forelimb and restoration of bipedal support during the stance phase, a postural feature that remained absent in vehicle-treated animals (Supplementary Fig. 8e-g).

## Discussion

We and others have shown that several AGC kinases constrain neurite outgrowth in vitro and axon regeneration in vivo^9,21,37–42^. Functional crosstalk among these kinases, rooted in their shared evolutionary origins, gives rise to biological robustness. This helps explain why single-target strategies have struggled to yield CNS regeneration therapies. Once recognized, this architecture becomes addressable by either drug combinations or single-agent polypharmacology. Single-agent polypharmacology offers regulatory, pharmacokinetic, and toxicological advantages over combinations^43^, and is therefore preferable where target biology permits it. The shared ancestry of these kinases, which we organize into clusters belonging to 4 evolutionary clades, underlies the structural homology that allows co-engagement by a single small molecule.

The current work identifies a higher-level pattern that idTRAX was not designed to detect: compounds with the largest effects on neurite outgrowth simultaneously engage multiple targets in different clades. Prior work established RO48’s in vivo efficacy across multiple mouse SCI models^9,14^. TMP-316 was developed from RO48 through phenotype-guided optimization that preserved multi-target pharmacology while meeting the clinical target product profile (Extended Data Table 2). The observation that RO48’s polypharmacology was retained in TMP-316 under phenotypic SAR pressure, without being selected for directly, is consistent with the multitarget activity being required for maximal neurite outgrowth promotion. Direct causal contributions to neurite outgrowth in vitro were demonstrated for one representative from each of the four clusters: ROCK2, S6K1, PKCγ, and PKX. These kinases were selected as likely primary drivers within their respective clusters; whether other cluster members contribute independently remains an open question. PKX’s role as a regulator of neurite outgrowth is unexplored; its phylogenetic and structural proximity to PKA suggests a role in cAMP-linked regulation. PKX’s substrate repertoire, downstream signaling, and neuronal roles remain to be characterized. In this study, potential PKX substrates were inferred from an atlas of kinase substrate peptides. The kinome-wide background pharmacological linkage between ROCK2 and PKX is among the highest reported in the literature (Fig. 2e), and Y-27632, the most widely used ROCK inhibitor, co-inhibits PKX (Supplementary Fig. 9). The body of evidence historically attributed to ROCK inhibition in regeneration may therefore reflect unrecognized support for PKX as an additional regulator.

Several members of the identified clusters have been independently implicated in axon growth regulation and pursued as single targets: ROCK^37^, PKC^38,39^, PKA^40^, and PKG^41,42^; however, their functional interrelatedness in regulating neurite and axon growth had not been recognized. Many tool compounds used to establish these targets, such as Y-27632 and Gö 6976, engage multiple AGC kinases (Supplementary Fig. 9). Preclinical efficacy, however, has been traditionally attributed to each compound’s nominal target (ROCK for Y-27632, PKC for Gö 6976)^20,38,39^. Within the same study, C3 transferase (selective for Rho/ROCK) failed to promote corticospinal sprouting or long-distance regeneration, while Y-27632 enhanced both^20^, suggesting that Y-27632’s polypharmacology likely contributed to its efficacy.

Compensatory coverage among the four clusters provides a molecular basis for the polypharmacology requirement, whereby combined inhibition both overcomes redundancy and recruits non-redundant activity ^6,7,44^. Phosphoproteomic profiling shows that combining S6K1i and ROCKi does not phenocopy RO48 or TMP-316 at the pathway or phosphosite level. This divergence indicates that TMP-316 polypharmacology engages multiple MoAs. These include reduced PTEN abundance along with PI3K hyperactivation; reorganization of the microtubule and actin cytoskeleton; growth-cone remodeling through Rac1/Cdc42 effectors; and PKC-dependent signaling. Strong representation of vesicle-trafficking pathways and MARCKS dephosphorylation suggest a role for secretory regulation, consistent with evidence that reduced presynaptic vesicle priming promotes CNS axon regeneration^45^. Direct S6K1 inhibition also decouples PI3K induction from downstream suppression of autophagy, allowing concomitant autophagy induction to further support axon growth^46^ (Fig. 6). Compartmentalization experiments show that simultaneous somatic and axonal application of RO48 produced neurite outgrowth exceeding compartment-restricted applications. This bolsters the rationale for lumbar IT delivery, which exposes both spinal axons and brain-resident somata. SAR concordance between hiGlut and prHP neurons, together with sequence and expression conservation across species, supports clinical translatability to humans.

**Figure 6.**
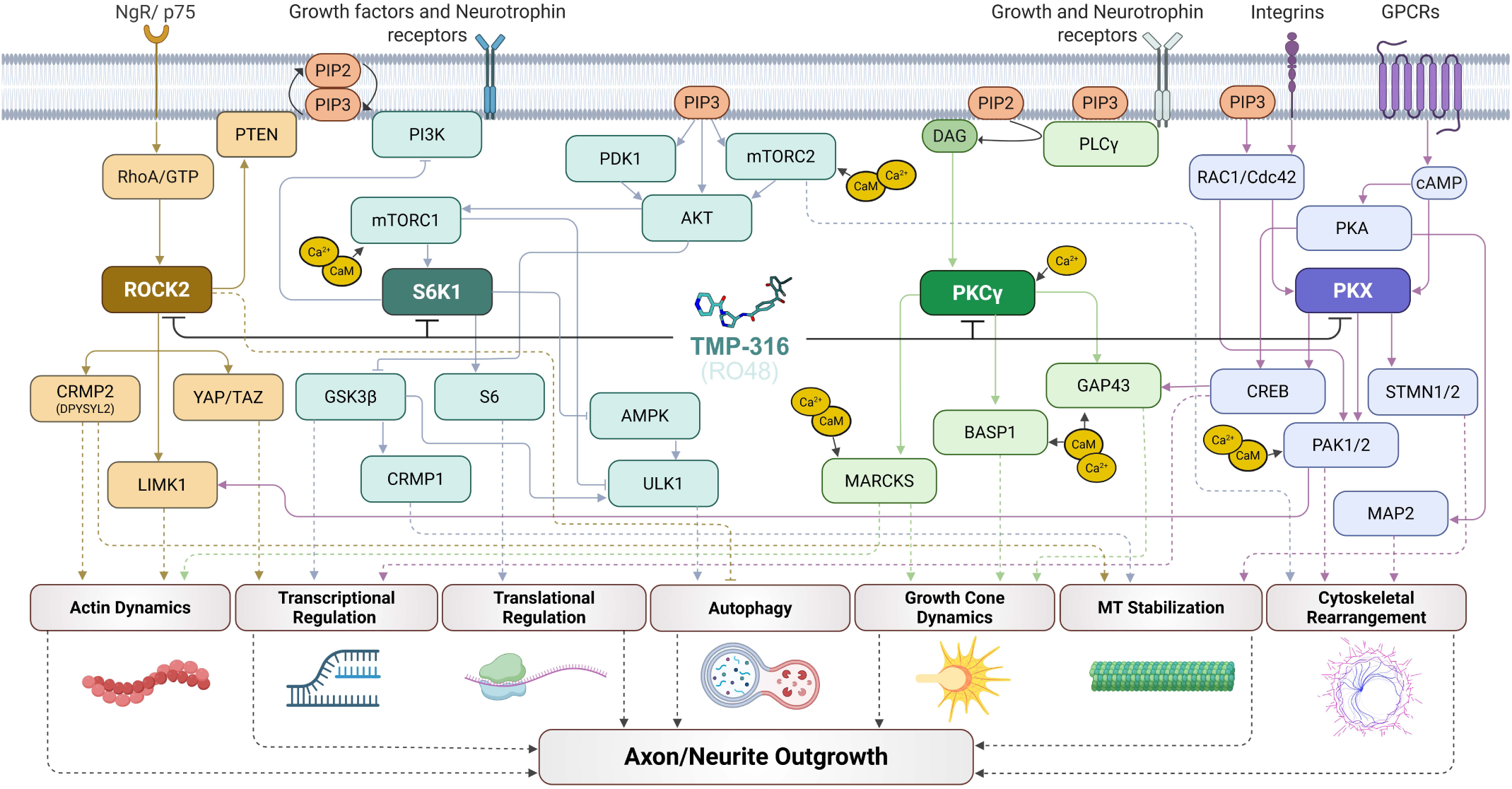
| Proposed multi-target mechanism of RO48 and TMP-316. Network polypharmacology diagram illustrating the signaling pathways downstream of simultaneous inhibition of ROCK2, S6K1, PKCγ, and PKX by RO48 and TMP-316. Target kinases are shown in dark fill. Solid arrows indicate activation; blunt arrows indicate inhibition; dashed arrows indicate indirect or multi-step relationships. Created in BioRender. Badillo-Martinez, A. (2026) https://BioRender.com/jvteqvy.

Prior work on RO48 demonstrated extensive histological evidence for axon regeneration, sprouting, and functional recovery in multiple mouse SCI models^9,14^. The current animal study asked whether TMP-316, delivered via the intended clinical route, could produce functional recovery in the rat cervical hemicontusion model ^35^, which recapitulates the injury level and forelimb functional deficits most prevalent in the clinical SCI population^36^. A single IT dose produced sustained motor recovery and restoration of weight-bearing behavior on the injured forelimb through the 42-day endpoint, which encompasses the post-acute window for this model^35^. This observation is consistent with the demonstration that RO48 upregulates a variety of regeneration-associated transcription factors in cultured neurons^47^. Given that post-injury edema confounds IT delivery and requires dedicated empirical optimization, dosing at delayed timepoints was deferred to IND-enabling work.

This work demonstrates an effective strategy for promoting CNS regeneration, whereby coordinated inhibition of multiple AGC kinases, achievable in a single small molecule, produces sustained functional recovery and defines a pharmacological profile that future candidates and modalities can target. It also proposes a discovery principle: when redundancy or overlap among evolutionarily related targets is identified prospectively, it simultaneously specifies the required engagement profile and makes single-agent polypharmacology structurally achievable. Future programs can apply platforms such as idTRAX with functional and evolutionary analyses to identify such target sets and design compounds for the required profile from the outset, rather than recovering polypharmacology retrospectively from phenotypic hits. AI-enabled generative chemistry conditioned on multi-target objectives^48^, together with high-throughput engagement profiling in live cells, makes this approach tractable for diseases governed by functionally coordinated targets.

## Methods

### Small molecule kinase inhibitors

RO48 and Blueprint Neurotherapeutics Network (BPN) program analogs were synthesized by Curia (Albany, NY) and assigned unique BPN identifiers, e.g., BPN-0037316 (TMP-316), BPN-0034858 (RO48). Earlier program analogs were synthesized at Sanford Burnham Prebys Medical Discovery Institute (SBI, Lake Nona, FL) and assigned unique SBI identifiers. Compounds were dissolved in DMSO for in vitro use. Additional details on small molecule compounds and the clinical formulation of TMP-316 are provided in Supplementary Methods.

### Neuronal cultures and neurite outgrowth assays

Rat primary hippocampal neurons and human iPSC-derived glutamatergic neurons (hiGluts; ioGlutamatergic Neurons, bit.bio) were cultured as previously described^18^. For neurite outgrowth assays, neurons were seeded onto PDL-coated 96-well plates (2,000 rat or 4,000 hiGlut cells/well), treated with compounds for 48 h, then fixed, immunostained for βIII-tubulin (Sigma-Aldrich, T2200, RRID: AB_262133, 1:1000) and Hoechst (nuclear DNA). Plates were imaged on an Opera Phenix High Content Screening (HCS) system, and neurite total length (NTL) was quantified in Harmony software. NTL was normalized to DMSO vehicle controls within each plate (%NTL), and a benchmarked %NTL was computed relative to the RO48 reference within each run. The drug effect score (DES), defined as the area above baseline under the dose-response curve, was normalized to the RO48 DES within the same assay run to yield %DES. Full cell culture, staining, and benchmarking details are provided in Supplementary Methods.

### siRNA-mediated gene silencing

Gene silencing was performed using Accell SMARTpool siRNA (Horizon Discovery), which enables transfection reagent-free, efficient knockdown in primary neurons^9,49^. siRNAs targeting *PRKCG* (Gene ID 24681; E-096727-00-0020) and *PRKX* (Gene ID 501563; E-106558-00-0020) were used to silence PKCγ (si-α-PKCγ) and PKX (si-α-PKX), respectively. An equimolar mixture of non-targeting siRNAs (D-001910-01-20, D-001910-02-20, and D-001910-04-20) served as a negative control (scramble).

E18 rat hippocampal neurons were seeded in 96-well plates at 1,800 cells/well in 100 µL NbActiv4 and incubated overnight at 37°C with 5% CO_2_. At 1 day *in vitro* (DIV), 50 µL of a 3 µM working solution of siRNA prepared in NbActiv4 was added directly to the medium, yielding a final concentration of 1 µM. After 24 h siRNA incubation, cells were treated with RO48, S6K1i, or ROCKi for 48 h, then fixed, stained, and analyzed using the Phenix HCS system as described above. Knockdown was verified by RT-qPCR in parallel cultures at 48 h post-siRNA addition, confirming target silencing during this inhibitor exposure (Supplementary Methods). NTL was normalized to the scramble control within each plate and expressed as robust Z-score according to the formula:

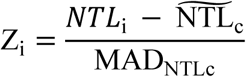

Where NTL_i_ is the value for the test well, and 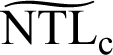 and MAD_NTLc_ are the median and median absolute deviation of control wells, respectively.

### Microfluidic axotomy assay

Neuronal soma and axons were isolated using two-compartment microfluidic devices (XonaChip 450 µm barrier, Xona Microfluidics) as previously described^50^. E18 hippocampal neurons were seeded into the somatic compartment and, at 8 DIV, axotomized by vacuum aspiration of the axonal compartment. Medium in the somatic compartment, the axonal compartment, or both was replaced with NbActiv4 containing RO48 [2 µM] or DMSO vehicle. After 48 h, neurons were fixed and immunostained for βIII-tubulin, imaged on a Dragonfly spinning-disk confocal microscope. Axon regrowth was quantified from maximum-intensity projections using an automated MATLAB (R2025b) tracing script modified from Wu et al.^51^, with values converted to robust z-scores. Full device, treatment, and imaging details are provided in Supplementary Methods.

### In-cell target engagement

In-cell target engagement of S6K1, ROCK2, PKCγ, and PKX was assessed using the NanoBRET Target Engagement Assay (Promega). HEK293T cells were resuspended in Opti-MEM I (Life Technologies) with 1% FBS (Cytiva) at 2 × 10^5^ cells/mL, transfected with kinase-NanoLuc fusion constructs (Promega) using FuGENE HD (Promega), and seeded at 20,000 cells/well in 100 µL Opti-MEM in white-wall 96-well plates. Each run included two technical replicates. Following overnight expression, NanoBRET tracers (K-9 for ROCK2; K-10 for PKX, S6K1, and PKCγ) were added at a final concentration of 0.5 µM, followed by test compounds at serial 1:10 concentrations ranging from 10^-8^ to 10 µM. For PKCγ assays, cells were pre-treated with 1 nM phorbol 12-myristate 13-acetate to activate the kinase prior to compound and tracer addition. Plates were incubated for 2 h at 37 °C with 5% CO₂, equilibrated at room temperature for 15 min, NanoBRET Nano-Glo substrate and Extracellular NanoLuc Inhibitor were added according to manufacturer instructions, and BRET signal was measured on a GloMax Discover System (Promega).

A four-parameter logistic model was fitted to the normalized data in GraphPad Prism (v11.0) to estimate the *K_d_apparent_* (*K_dapp_*) for each compound–target pair. Percent target engagement (%TE) at concentrations with benchmarked %NTL values was calculated according to:

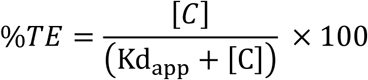

where [*C*] is the compound concentration in µM.

### Multi-target modeling

To assess the relationship between multi-target engagement and neurite outgrowth activity, the arithmetic sum of %TE values for S6K1, ROCK2, PKCγ, and PKX, with a maximum of 400%, was calculated at each concentration tested in the neurite outgrowth assay. Benchmarked %NTL was plotted as a function of %TE sum across all compounds and concentrations, and a moving average was superimposed to visualize the overall trend between aggregate target engagement and neurite outgrowth activity.

### *T_ij_* Analysis

Pairwise co-engagement was quantified using the activity-profile Tanimoto coefficient (*T_ij_*) as defined by Metz et al. (Nat Chem Biol 2011, 7:200). Per-observation target engagement (%TE) values for the four kinases (S6K1, ROCK2, PKX, PKCγ) were binarized as engaged at %TE ≥ 25%. For each of the six kinase pairs, *T_ij_* was computed as |A ∩ B| / |A ∪ B|, where A and B are the sets of observations engaged on each kinase of the pair. To test whether co-engagement scales with cell effect, the 711 observations, pooled across all compounds and concentrations, were stratified into equal-count quintiles by Benchmarked NTL (Q1: −4.67 to 2.42; Q2: 2.45 to 5.38; Q3: 5.38 to 10.26; Q4: 10.27 to 29.95; Q5: 30.29 to 174.86; n = 142–143 per quintile), and *T_ij_* was calculated within each stratum. To ensure *T_ij_* estimates were supported by a sufficient number of compounds engaging at least one kinase in the pair, *T_ij_* was not calculated when |A ∪ B| < 10. Background *T_ij_* values for the same six kinase pairs were taken directly from Supplementary Table 2 of Metz et al.^52^ (3,858 compounds × 172 kinases, Abbott internal kinase profiling panel), providing a kinome-wide reference. Both per-pair *T_ij_* and the cross-pair mean per stratum were compared against this background and against the Metz et al threshold of *T_ij_* ≥ 0.55 for a strong pharmacological relationship.

### Western blot of cell lysates

prHP neurons were treated with TMP-316 across a serial concentration range (0.8 nM–10 µM), or with vehicle (DMSO), S6K1i (PF-4708671, 5 µM), ROCKi (Rho Kinase inhibitor IV, 10 µM), S6K1i + ROCKi, RO48 (2 µM), or TMP-316 (2 µM), and analyzed by western blot. Cells were harvested at 1h to capture acute phosphorylation responses, or at 24h to allow sufficient time for changes in protein abundance. Cells were washed once in phosphate buffer saline (PBS) and then lysed in hot 4% SDS loading buffer as previously described^21^. Proteins were separated on NuPAGE Bis-Tris Mini Protein Gels, 4–12%, 1.0–1.5 mm (ThermoFisher) in NuPAGE MOPS SDS Running Buffer (ThermoFisher) and transferred to nitrocellulose membranes (0.45 μm, Millipore) in NuPage Transfer Buffer (ThermoFisher). Membranes were probed with primary antibodies against phospho-S6 Ser240/244 (CST 5364L, RRID:AB_10694233), total S6 (CST 2317S, RRID:AB_2238583), phospho-Akt Ser473 (CST 4058, RRID:AB_331168), pan-Akt (CST 2920, RRID:AB_1147620), PTEN (CST 9559T, RRID:AB_390810), or β-actin (Sigma A5441, RRID:AB_476744) at manufacturer recommended dilutions. Proteins were visualized using secondary antibodies (680 RD or 800 CW, Licor) and imaged on an Azure C600 instrument. Band intensity was quantified using AzureSpot Pro software.

### Phosphoproteomic analysis

Quantitative phosphoproteomic profiling was performed in primary E18 rat hippocampal neurons treated with TMP-316 [2 µM], RO48 [2 µM], ROCKi [10 µM], S6K1i [5 µM], or ROCKi+S6K1i for 2 or 24 h, alongside vehicle controls, in three biological replicates. These concentrations were selected as those that maximized neurite outgrowth without causing cellular toxicity (>90% viability relative to vehicle; Supplementary Fig. 1k). Following lysis, digestion, and sequential metal-oxide/Fe-NTA (SMOAC) phosphopeptide enrichment, samples were analyzed by data-independent acquisition on a Thermo Orbitrap Eclipse mass spectrometer. Peptides were identified and quantified in Proteome Discoverer 3.2 using CHIMERYS against the rat reference proteome (UniProt UP000002494) at a 1% false discovery rate, with label-free quantification of phosphopeptides. Differential phosphorylation was assessed relative to time-matched vehicle controls. Full sample-processing and search parameters are provided in Supplementary Methods.

### Pathway analysis

Differentially phosphorylated protein lists were generated for each treatment relative to time-matched vehicle by filtering the protein-level quantification (Supplementary Table 5) to phosphorylation-annotated entries with |log2 fold change| > 0.5 and adjusted p < 0.1. Retained proteins were mapped to gene symbols and submitted to Metascape (v3.5.20250701) for GO Biological Process enrichment against the Rattus norvegicus whole genome, with terms of enrichment factor > 1.5 and LogP < −2 retained (Supplementary Data Tables 6 and 7).

Significant terms were partitioned into mutually exclusive biological categories and summarized per category-condition pair using a size-corrected composite score. Full thresholds and category-scoring details are provided in Supplementary Methods.

### Computational docking and molecular dynamics simulation

Protein-ligand complexes were predicted by co-folding in Boltz-2 (v2.2.0)^53^, using two structural templates per complex: the full-length target from the AlphaFold Protein Structure Database and the PKA catalytic subunit in complex with RO48 (PDB 1SVG). Complexes were refined by a simulated annealing routine in OpenMM (v8.4.0). Structural representations were generated in PyMOL (Schrödinger). Full co-folding parameters and the molecular dynamics pose-optimization protocol are provided in Supplementary Methods.

### Substrate preference analysis

Percentile substrate scores for AGC kinases were extracted from published data profiling over 89,000 substrate peptides^29^. Pearson correlation coefficients were computed in MATLAB (R2025b) across all substrates for all profiled AGC kinases. Hierarchical clustering was performed on a dissimilarity matrix defined as 1 − ρ, where ρ is the Pearson correlation coefficient, using the unweighted pair group method with arithmetic mean.

### Animals

All animal procedures were approved by the Institutional Animal Care and Use Committee at the University of Miami Miller School of Medicine and performed in accordance with the National Institutes of Health Guide for the Care and Use of Laboratory Animals. Animals were housed under standard conditions with ad libitum access to food and water.

### Surgical SCI and drug delivery procedures

Sprague-Dawley male rats of uniform age and weight range were selected for this study for two reasons. First, graded cervical hemicontusion model was characterized in adult male rats^35^. Second, body mass variability directly affects intrathecal dose per kilogram and would introduce a confound orthogonal to the focused question this study was designed to address, whether a single IT dose of TMP-316 produces functional recovery in a clinically relevant cervical hemicontusion model. Rats (7 weeks old) were randomly assigned to three experimental groups (n = 12–13 per group): vehicle control, TMP-316 at ∼0.3 mg/kg, and TMP-316 at ∼1 mg/kg. TMP-316 was administered intrathecally directly prior to injury, with anesthesia maintained. This timing was selected because post-injury edema could impede rostral drug distribution from the lumbar intrathecal space. IT catheterization and drug delivery were performed at the L6–S1 interlaminar space as previously described by Elwardany et al^54^. To apply the SCI injury, animals were anesthetized (ketamine 75 mg/kg i.p., xylazine 5 mg/kg i.p., maintained with isofluorane), given an analgesic (Buprenorphine SR 0.58 mg/kg subQ), and placed in a prone position on a heated surgical platform maintained at 37 °C.. The surgical site was shaved and sterilized, and a midline dorsal incision was made over the cervical spine. The paraspinal musculature was reflected bilaterally to expose the C5–C6 vertebral laminae. A laminectomy was performed at the C5–C6 level to expose the dorsal surface of the spinal cord, leaving the dura intact. The right half of the spinal cord was injured using the Infinite Horizon impactor set to a target force of 170 kilodynes (kdyn); actual force and displacement values were recorded in real time. Personnel were blinded to treatment allocation. Following injury, the musculature was sutured in layers, and the skin was closed. Bladders were manually expressed twice a day until micturition fully recovered. Predefined exclusion criteria were applied: drop below 80% of pre-surgical body weight, loss of movement of both hindlimbs after surgery, or premature sacrifice for morbidity; per-animal inclusion/exclusion and mortality are reported at the Open Data Commons for Spinal Cord Injury^55^ (ODC-SCI; RRID:SCR_016673) and are available for download.

### Behavioral assessment

Locomotor recovery was assessed using kinematic gait analysis with the MotoRater system (TSE Systems, Bad Homburg, Germany)^56^. Animals were habituated and trained on the overground walking runway for one week prior to injury; the final session served as the pre-injury baseline. Post-injury recordings were conducted weekly for 7 – 42 days post-injury (DPI), with five trials acquired per animal per session. Behavioral assessments were performed fully blinded to treatment.

### Analysis of kinematic data

Kinematic analyses were performed in Python. MotoRater time-series data (46 base parameters per trial) were processed with automated feature extraction (TSFresh v0.21.1)^57^, followed by a selection procedure identifying features responsive to injury and to treatment, yielding 44 features from 9 base parameters. Recovery was quantified using a composite kinematic score (CKS), computed by principal component analysis of the selected features and Euclidean projection onto the baseline-to-acute-injury axis, normalized to a 0-100 scale (0 = acutely injured vehicle at 7 DPI; 100 = baseline at 0 DPI). Three additional, mathematically distinct recovery metrics were computed to verify method independence. Full feature-extraction, selection, and recovery-metric details are provided in Supplementary Methods.

### Kinematic trace analysis

Forelimb gait cycle was characterized using toe-to-ear traces, computed by expressing the right forelimb toe position relative to the right ear for each video frame, yielding a body-centered trajectory independent of forward translation. Traces were cycle-segmented, averaged across trials and animals to yield group-level consensus traces ± SEM, and circularly normalized for cross-group comparison. Full computational details are provided in Supplementary Methods.

### Quantification of spinal cord lesion size

At 42 DPI, animals were perfused and spinal cords post-fixed, then a segment centered on the injury epicenter was sectioned in the transverse plane. Sections were immunostained for GFAP (Agilent/Dako, Z0334, RRID: AB_10013382) with Hoechst counterstain and imaged on a Dragonfly spinning-disk confocal microscope. Histological images were analyzed using a custom deep learning-assisted pipeline (U-Net) to segment the spinal cord boundary and classify tissue as injured, uninjured, or lesion. Lesion size (%) was calculated per section as lesion area divided by total spinal cord cross-sectional area, averaged across sections per animal, and summarized as median values with 95% confidence intervals. Full perfusion, sectioning, staining, imaging, and analysis details are provided in Supplementary Methods.

### In vitro profiling assays

Biochemical binding affinity (Kd) was determined at Eurofins DiscoverX Corporation (San Diego, CA, USA) using the KINOMEscan™ KdELECT active-site competition binding platform. Kinase inhibition profiling of TMP-316 was performed by Reaction Biology Corporation (Malvern, PA, USA) against a panel of 353 wild-type kinases at a single compound concentration of 500 nM in the presence of 1 mM adenosine triphosphate (ATP), in duplicate; results were expressed as percent enzyme activity relative to dimethyl sulfoxide (DMSO) controls, with staurosporine as the reference inhibitor. RO48 was profiled previously on the same platform in a similar manner but using a concentration of 100 nM in the presence of Km ATP, against a panel of 251 wild-type kinases; these data are provided for reference. Given the differing assay conditions, inhibition magnitudes were not compared between compounds. Cross-compound comparison was limited to kinases inhibited by RO48 and showing no detectable inhibition by TMP-316. Kinase activities were converted into percent inhibition and mapped onto the human kinome tree using KinMap^58^. hERG channel (Kv11.1) inhibition was assessed by automated whole-cell patch clamp (QPatch) in HEK293 cells stably expressing human KCNH2, at physiologic temperature, across six concentrations ranging from 0.10 to 30 µM; verapamil served as a positive control (IC50 = 420 nM). P-glycoprotein (Pgp) efflux was determined using the MDR1-MDCK Permeability Assay at Curia (Albany, NY, USA). The bacterial reverse mutation (Ames) assay was performed by Frontage Laboratories under GLP conditions using Salmonella typhimurium strains TA98, TA100, TA1535, and TA1537 and Escherichia coli strain WP2uvrA, in the presence and absence of an exogenous mammalian metabolic activation system (S9), across seven concentrations ranging from 5 to 5,000 µg/plate.

### Gini coefficient calculation

The selectivity of TMP-316 across the kinome was quantified using the Gini coefficient, calculated from percent inhibition obtained from kinome profiling data. Prior to analysis, values below 0% and above 100% were clipped to 0% and 100%, respectively. Kinases returning no value were excluded. The Gini coefficient was computed according to Graczyk^59^ as:

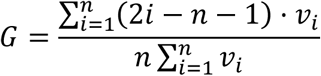

where v₁ ≤ v₂ ≤ … ≤ vₙ are the rank-ordered percent inhibition values across n kinases. A Gini coefficient of 0 indicates equal inhibition across all kinases (no selectivity); a value of 1 indicates complete selectivity for a single kinase. Results were verified by computing 1 − 2 × area under the Lorenz curve using the trapezoidal rule, confirming numerical equivalence with the closed-form calculation. All analyses were performed in MATLAB (R2025b).

## Statistical analysis

Statistical analyses were performed in GraphPad Prism v11.0 and presented as mean ± SEM unless otherwise stated. Two-group comparisons were evaluated using an unpaired t-test. Comparisons across three or more independent groups were assessed using a one-way analysis of variance (ANOVA). Post-hoc analyses were conducted using Dunnett’s multiple comparison test when comparing groups to a single control or Tukey’s multiple comparison test when comparing all groups to one another. For longitudinal behavioral data, group differences across time points were analyzed using Repeated Measures ANOVA with Dunnett’s multiple comparison test. Statistical significance was defined as p < 0.05.

## Acknowledgements

The authors thank Sierra Jade Bagwell, Aya Hanna, Christian Ebo, Jennifer Maria Campos, Liza Hourany, Rachaeel Martinez, Mia Begera, Nicolas Romero, Camila Banciella, Hailey Weintraub, Erik Fernandez, Ally Gallagher, Benjamin McCulley, Seth Herr, Brian Kang, and Indigo Williams for technical support and assistance in data acquisition; Matthew Scarmuzza, Kevin Ma, Javier Carrillo, Hannah Beatty, Emily Owens, Katarzyna Jagieslska, and Joey Gawler for analytical support; Mahesh Gokara and The Miami Project Drug Discovery Core (RRID:SCR_022542) for data acquisition and processing; Yan Shi and The Miami Project Imaging Core Facility (RRID:SCR_025492) for imaging support; Maria Torres and The Miami Project Histology Shared Resource for histological support; Ramon A. German of the University of Miami Division of Veterinary Resources at the Lois Pope LIFE Center for assistance with surgeries and animal care; James Vasta and Matthew Robers at Promega Corporation for technical guidance on NanoBRET Target Engagement assays; and LeeAnn Higgins and Todd Markowski at the Center for Metabolomics and Proteomics, University of Minnesota, for proteomics data generation and support. During this work, Omar Elwardany was on research leave from the Department of Neurosurgery, Faculty of Medicine, Suez Canal University, Ismailia, Egypt. The authors acknowledge the officials and consultants of the National Institutes of Health Blueprint Neurotherapeutics Network for scientific and programmatic guidance.

## Author Contributions

HA, ABM, BA, JLB, VPL, JKL, and GJ designed experiments; HA, ABM, BA, BBS, SS, RL, OSE, VPL, JKL and RB performed experiments; HA, ABM, BA, BBS, SS, JLB, NG, JKL and PS analyzed data; HA, ABM, BA, and BBS wrote the manuscript; HA, VPL, JLB and JKL acquired funding and supervised the study; all authors reviewed and edited the manuscript.

## Supplementary Information

This file contains associated Supplementary Material included with the initial submission.

## Extended Data Tables

**Extended Data Table 1:** AGC Kinases Identified by idTRAX as Potential Repressors of Neurite Outgrowth.

| Protein name | Common protein name | Official gene name | UniProt-annotated functions relevant to neuronal growth regulation<sup>1</sup> | %Identity Human/Rat (Total Sequence)<sup>2</sup> | %Identity Human/Rat (Kinase Domain)<sup>2</sup> | Neuronal expression<sup>3</sup> | Somatodendritic protein evidence (PMID)<sup>4</sup> | Axonal protein evidence (PMID)<sup>4</sup> |
| --- | --- | --- | --- | --- | --- | --- | --- | --- |
| S6K1 | Ribosomal protein S6 kinase beta-1 | RPS6KB1 | Effector of mTORC1; regulates translation initiation and elongation through phosphorylation of RPS6, EIF4B, PDCD4 and EEF2K; suppresses autophagy within the PI3K pathway; participates in feedback inhibition of PI3K/mTORC2/AKT signaling. Activated by mTORC1 (Thr-412) or PDPK1 (Thr-252). | 99.6% | 100.0% | Yes | 14623952 | 35098924 |
| S6K2 | Ribosomal protein S6 kinase beta-2 | RPS6KB2 | Acts downstream of mTOR in response to growth factors and nutrients to regulate translation via phosphorylation of ribosomal protein S6; component of PI3K/mTOR-linked translational control. | 93.7% | 95.8% | Yes |  |  |
| RSK1 | Ribosomal protein S6 kinase alpha-1 | RPS6KA1 | Acts downstream of ERK; regulates translation through phosphorylation of RPS6 and EIF4B and modulates mTOR signaling via TSC2 and RPTOR; mediates cell survival by repressing the pro-apoptotic proteins BAD and DAPK1; activates transcription factors including CREB1. | 97.6% | 98.9% | Yes | 15893597, 35648763 |  |
| RSK3 | Ribosomal protein S6 kinase alpha-2 | RPS6KA2 | Acts downstream of ERK signaling; regulates translation and modulates mTOR-linked growth signaling; relays mitogenic and stress signals to transcription-factor activation; contributes to cell survival. | 95.6% | 98.5% | Yes | 37068088 | 36239373 |
| RSK2 | Ribosomal protein S6 kinase alpha-3 | RPS6KA3 | Acts downstream of ERK; regulates translation through RPS6 and EIF4B phosphorylation and modulates mTOR signaling; represses the pro-apoptotic proteins BAD and DAPK1; propagates mitogenic, stress, insulin and growth-factor signals. | 99.7% | 100.0% | Yes | 37068088 | 36239373 |
| RSK4 | Ribosomal protein S6 kinase alpha-6 | RPS6KA6 | Growth-factor-independent kinase; participates in p53/TP53-dependent cell-growth-arrest signaling and plays an inhibitory role during embryogenesis. | 88.8% | 98.1% | Yes |  | 36239373 |
| MSK1 | Ribosomal protein S6 kinase alpha-5 | RPS6KA5 | Acts downstream of ERK1/2 and p38 MAPK; phosphorylates and activates the transcription factors CREB1 and ATF1 and phosphorylates histone H3, coupling mitogenic and stress signals to gene expression. | 96.6% | 97.0% | Yes | 15893597 | 36239373 |
| MSK2 | Ribosomal protein S6 kinase alpha-4 | RPS6KA4 | Acts downstream of ERK1/2 and p38 MAPK to phosphorylate CREB1, ATF1 and RELA and to phosphorylate histone H3, mediating mitogen- and stress-induced transcriptional activation. | 96.4% | 98.1% | Yes | 27376065 |  |
| ROCK1 | Rho-associated protein kinase 1 | ROCK1 | Key regulator of the actin cytoskeleton and cell polarity; controls stress-fiber and focal-adhesion formation, cell adhesion, motility and neurite retraction through substrates including LIMK1/2, MYL9/MLC2 and PPP1R12A. | 95.9% | 100.0% | Yes | 28396402 | 36239373 |
| ROCK2 | Rho-associated protein kinase 2 | ROCK2 | Regulator of the actin cytoskeleton and cell polarity; mediates neurite retraction and growth-cone dynamics through LIMK1/2 and the growth-cone regulator DPYSL2/CRMP2; predominant CNS isoform; also regulates spine and synaptic properties in the hippocampus. | 96.4% | 97.0% | Yes | 24832597, 28396402, 35098924 | 36239373, 35098924 |
| PKCα | Protein kinase C alpha type | PRKCA | Calcium- and DAG-activated kinase; regulates actin cytoskeletal dynamics and RhoA activation via RhoGDI phosphorylation; broad roles in proliferation, survival, migration and adhesion through direct substrates and ERK signaling. | 99.3% | 99.6% | Yes | 1303731, 35098924 | 7644028, 7897445, 36239373, 35098924 |
| PKCβ | Protein kinase C beta type | PRKCB | Calcium- and DAG-activated kinase involved in receptor signaling, ERK/MAPK activation and PI3K/AKT-linked regulation; contributes to survival and proliferation signaling. | 98.8% | 100.0% | Yes | 1303731, 35098924 | 7644028, 7897445, 36239373, 35098924 |
| PKCγ | Protein kinase C gamma type | PRKCG | Neuron-specific, calcium- and DAG-activated kinase; phosphorylates and regulates the neuronal receptors GRIA4/GLUR4 and GRIN1/NMDAR1; mediates synaptic function, synaptogenesis, synaptic plasticity, excitotoxicity and cell survival after ischemia. | 99.0% | 99.6% | Yes | 1303731, 35098924 | 7644028, 7897445, 36239373, 35098924 |
| PKX | -dependent protein kinase catalytic subunit 1 | PRKX | cAMP-dependent kinase acting through phosphorylation of downstream targets including CREB, SMAD6 and PKD1; functions in differentiation, epithelial morphogenesis, migration and proliferation. | 77.0% | 86.7% | Yes | 15879576 |  |
| PKG1 | cGMP-dependent protein kinase 1 | PRKG1 | Key mediator of the cGMP signaling pathway; phosphorylates targets regulating cellular calcium and CNS processes including axon guidance, hippocampal and cerebellar learning, circadian rhythm and nociception. | 98.1% | 99.2% | Yes | 30319353 | 30319353 |
| PKG2 | cGMP-dependent protein kinase 2 | PRKG2 | cGMP-dependent kinase; acts downstream of NMDAR to drive synaptic accumulation of GRIA1/GLUR1 and increase synaptic plasticity; activates ERK1/2 and regulates gene expression; also regulates intestinal secretion via CFTR and bone growth. | 96.3% | 98.8% | Yes | 11738129 | 11738129 |

<sup>1</sup>Information obtained from the UniProt Consortium , UniProt: the Universal Protein Knowledgebase in 2025, Nucleic Acids Research, Volume 53, Issue D1, 6 January 2025, Pages D609–D617, https://doi.org/10.1093/nar/gkac1010
<sup>2</sup>Determined by protein alignment using the UniProt database: https://www.uniprot.org/, accessed 06/2021
<sup>3</sup>Mathias Uhlén et al. Tissue-based map of the human proteome. Science. 347, 1260419 (2015). DOI:10.1126/science.1260419
<sup>4</sup>For somatodendritic and axonal protein evidence, blank cells mean that no qualifying protein-level compartment evidence was identified in the reviewed literature or datasets. Absence of evidence in these sources does not establish absence of the protein from that compartment.

**Extended Data Table 2:** TMP-316 Properties.

| Target Candidate Properties Profile | Optimal | Acceptable | TMP-316 |
| --- | --- | --- | --- |
| **Neurite Outgrowth** |  |  |  |
| Primary rat hippocampal (prHP) neurons |  |  |  |
| EC50 (μM) | ≤ 1 | ≤ 10 | 0.938 ± 0.185 (n = 13) |
| %DES | ≥ 90% | ≥ 50% | 134 ± 14.2% (n = 13) |
| Human iPSC-derived glutamatergic (hiGlut) |  |  |  |
| %DES | ≥ 90% | ≥ 50% | 97.3 ± 17.7 % (n = 2) |
| **In-Cell Target Engagement Assay** |  |  |  |
| S6K1 | apparent Kd ≤ 100 nM | apparent Kd ≤ 1 μM | 0.34 ± 0.37 μM (n=5) |
| ROCK2 | apparent Kd ≤ 100 nM | apparent Kd ≤ 1 μM | 0.039 ± 0.01 μM (n=3) |
| **Kinetic Solubility** |  |  |  |
| Artificial CSF (μM) | > 50 | > 10 | 82 |
| Formulation buffer (mM) | > 30 | > 20 | > 20 |
| **MDCK-MDR1** |  |  |  |
| Efflux ratio | < 3 | < 6 | 4.6 |
| **Preclinical Safety** |  |  |  |
| GLP Ames | Negative | Negative | Negative |
| hERG IC50 (μM) | > 30 | > 10 | > 30 |

**Extended Data Table 3:**
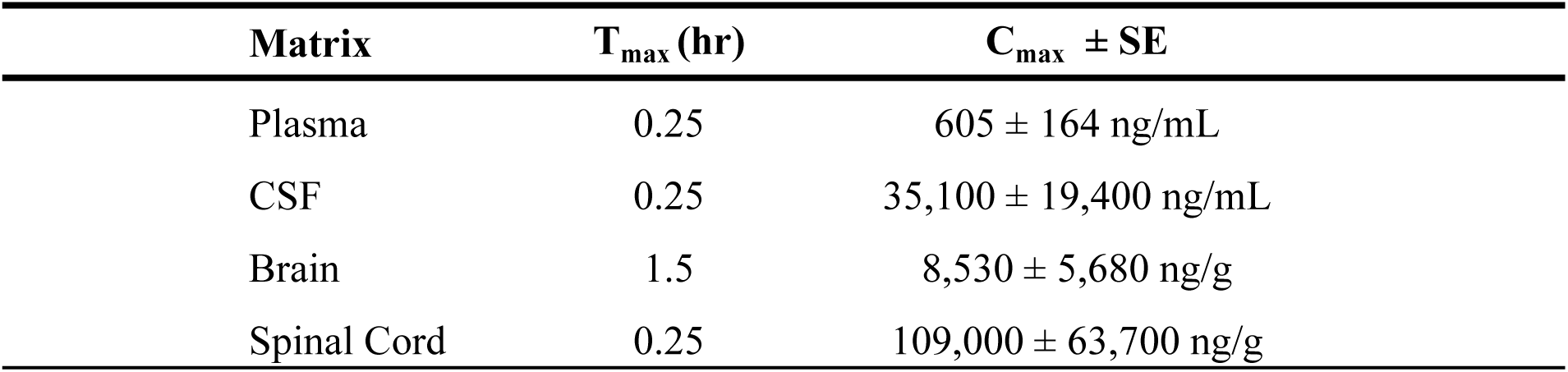
Pharmacokinetic Parameters of TMP-316 Following a Single ntrathecal Dose Administration to Male Sprague Dawley Rats.

**Extended Data Table 4:**
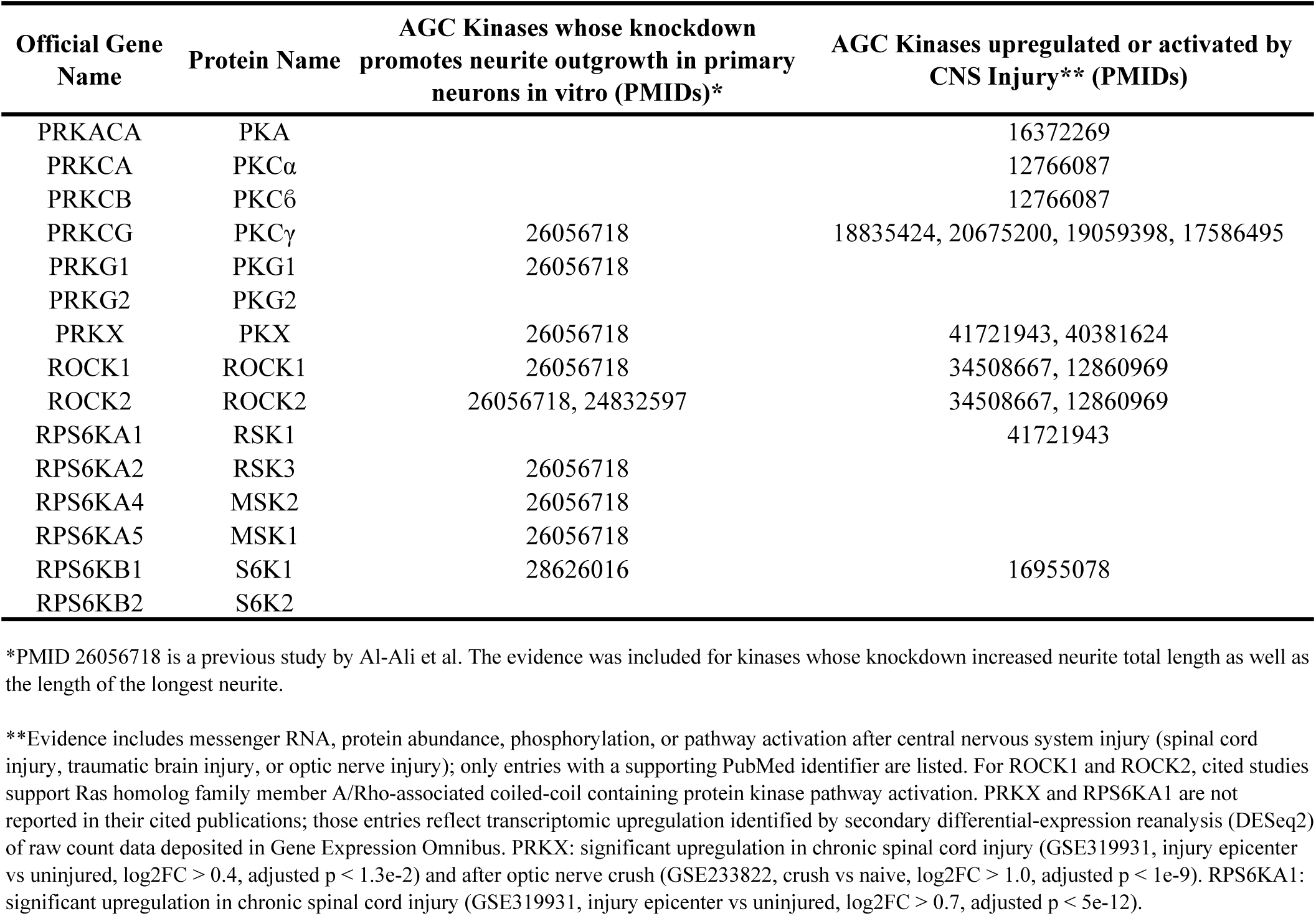
Genetic evidence of AGC kinase involvement in neurite outgrowth and CNS injury response.

**Extended Data Table 5:**
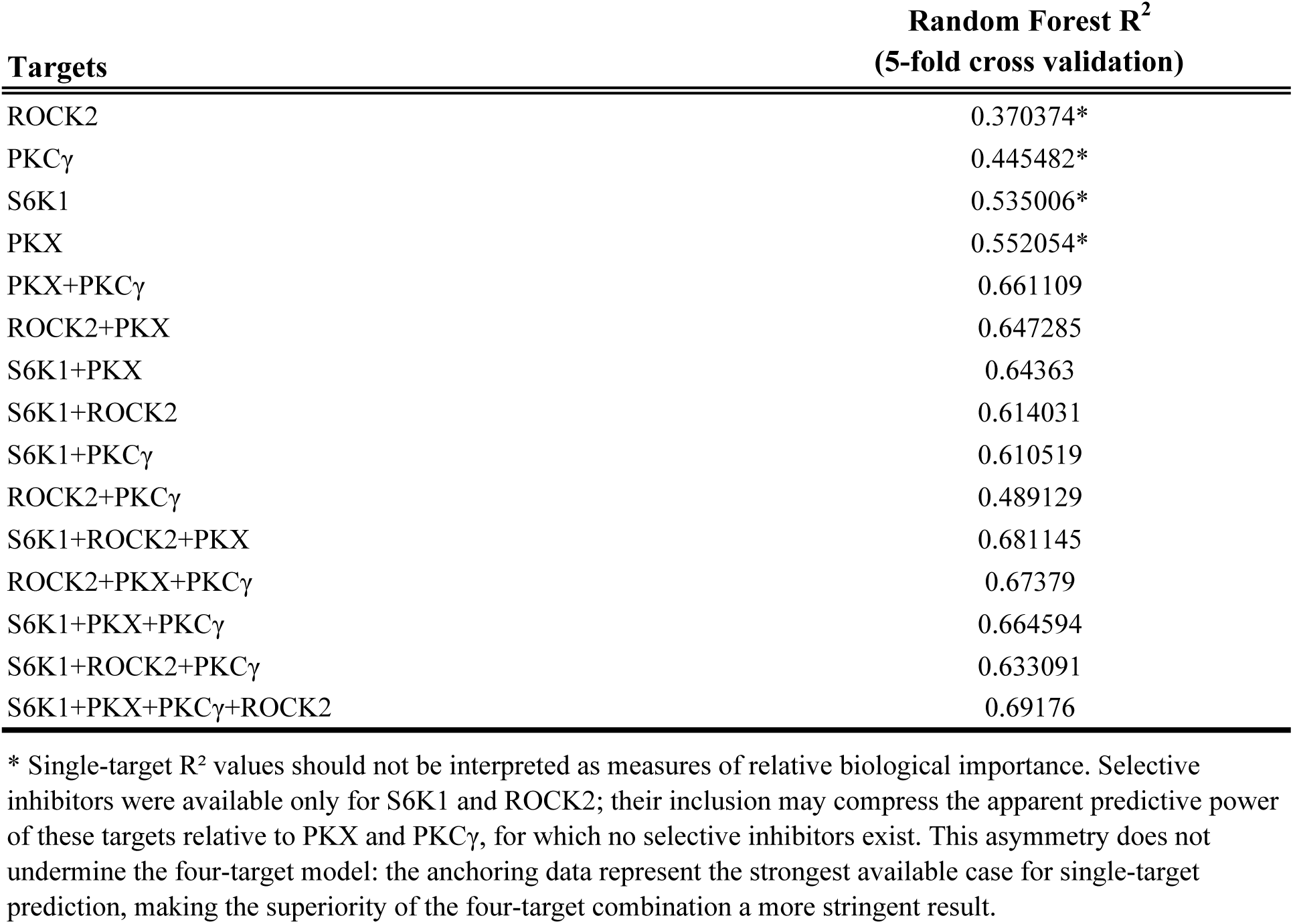
Random Forest Analysis of Target Combinations.

**Extended Data Table 6:**
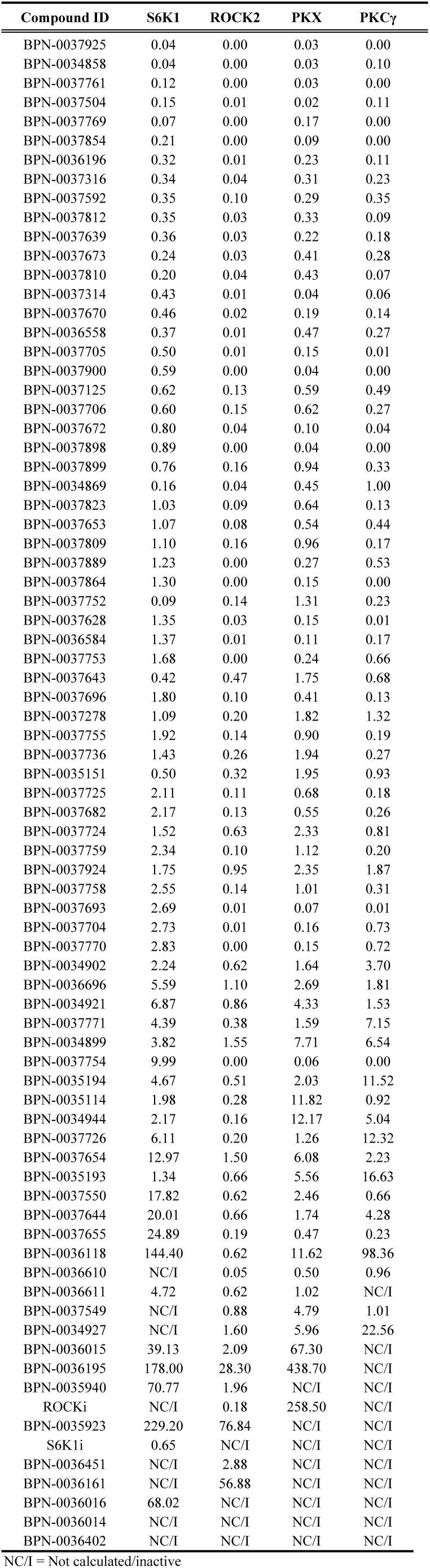
In-cell Apparent Kds (µM)

| Compound ID | S6K1 | ROCK2 | PKX | PKCγ |
| --- | --- | --- | --- | --- |
| BPN-0037925 | 0.04 | 0.00 | 0.03 | 0.00 |
| BPN-0034858 | 0.04 | 0.00 | 0.03 | 0.10 |
| BPN-0037761 | 0.12 | 0.00 | 0.03 | 0.00 |
| BPN-0037504 | 0.15 | 0.01 | 0.02 | 0.11 |
| BPN-0037769 | 0.07 | 0.00 | 0.17 | 0.00 |
| BPN-0037854 | 0.21 | 0.00 | 0.09 | 0.00 |
| BPN-0036196 | 0.32 | 0.01 | 0.23 | 0.11 |
| BPN-0037316 | 0.34 | 0.04 | 0.31 | 0.23 |
| BPN-0037592 | 0.35 | 0.10 | 0.29 | 0.35 |
| BPN-0037812 | 0.35 | 0.03 | 0.33 | 0.09 |
| BPN-0037639 | 0.36 | 0.03 | 0.22 | 0.18 |
| BPN-0037673 | 0.24 | 0.03 | 0.41 | 0.28 |
| BPN-0037810 | 0.20 | 0.04 | 0.43 | 0.07 |
| BPN-0037314 | 0.43 | 0.01 | 0.04 | 0.06 |
| BPN-0037670 | 0.46 | 0.02 | 0.19 | 0.14 |
| BPN-0036558 | 0.37 | 0.01 | 0.47 | 0.27 |
| BPN-0037705 | 0.50 | 0.01 | 0.15 | 0.01 |
| BPN-0037900 | 0.59 | 0.00 | 0.04 | 0.00 |
| BPN-0037125 | 0.62 | 0.13 | 0.59 | 0.49 |
| BPN-0037706 | 0.60 | 0.15 | 0.62 | 0.27 |
| BPN-0037672 | 0.80 | 0.04 | 0.10 | 0.04 |
| BPN-0037898 | 0.89 | 0.00 | 0.04 | 0.00 |
| BPN-0037899 | 0.76 | 0.16 | 0.94 | 0.33 |
| BPN-0034869 | 0.16 | 0.04 | 0.45 | 1.00 |
| BPN-0037823 | 1.03 | 0.09 | 0.64 | 0.13 |
| BPN-0037653 | 1.07 | 0.08 | 0.54 | 0.44 |
| BPN-0037809 | 1.10 | 0.16 | 0.96 | 0.17 |
| BPN-0037889 | 1.23 | 0.00 | 0.27 | 0.53 |
| BPN-0037864 | 1.30 | 0.00 | 0.15 | 0.00 |
| BPN-0037752 | 0.09 | 0.14 | 1.31 | 0.23 |
| BPN-0037628 | 1.35 | 0.03 | 0.15 | 0.01 |
| BPN-0036584 | 1.37 | 0.01 | 0.11 | 0.17 |
| BPN-0037753 | 1.68 | 0.00 | 0.24 | 0.66 |
| BPN-0037643 | 0.42 | 0.47 | 1.75 | 0.68 |
| BPN-0037696 | 1.80 | 0.10 | 0.41 | 0.13 |
| BPN-0037278 | 1.09 | 0.20 | 1.82 | 1.32 |
| BPN-0037755 | 1.92 | 0.14 | 0.90 | 0.19 |
| BPN-0037736 | 1.43 | 0.26 | 1.94 | 0.27 |
| BPN-0035151 | 0.50 | 0.32 | 1.95 | 0.93 |
| BPN-0037725 | 2.11 | 0.11 | 0.68 | 0.18 |
| BPN-0037682 | 2.17 | 0.13 | 0.55 | 0.26 |
| BPN-0037724 | 1.52 | 0.63 | 2.33 | 0.81 |
| BPN-0037759 | 2.34 | 0.10 | 1.12 | 0.20 |
| BPN-0037924 | 1.75 | 0.95 | 2.35 | 1.87 |
| BPN-0037758 | 2.55 | 0.14 | 1.01 | 0.31 |
| BPN-0037693 | 2.69 | 0.01 | 0.07 | 0.01 |
| BPN-0037704 | 2.73 | 0.01 | 0.16 | 0.73 |
| BPN-0037770 | 2.83 | 0.00 | 0.15 | 0.72 |
| BPN-0034902 | 2.24 | 0.62 | 1.64 | 3.70 |
| BPN-0036696 | 5.59 | 1.10 | 2.69 | 1.81 |
| BPN-0034921 | 6.87 | 0.86 | 4.33 | 1.53 |
| BPN-0037771 | 4.39 | 0.38 | 1.59 | 7.15 |
| BPN-0034899 | 3.82 | 1.55 | 7.71 | 6.54 |
| BPN-0037754 | 9.99 | 0.00 | 0.06 | 0.00 |
| BPN-0035194 | 4.67 | 0.51 | 2.03 | 11.52 |
| BPN-0035114 | 1.98 | 0.28 | 11.82 | 0.92 |
| BPN-0034944 | 2.17 | 0.16 | 12.17 | 5.04 |
| BPN-0037726 | 6.11 | 0.20 | 1.26 | 12.32 |
| BPN-0037654 | 12.97 | 1.50 | 6.08 | 2.23 |
| BPN-0035193 | 1.34 | 0.66 | 5.56 | 16.63 |
| BPN-0037550 | 17.82 | 0.62 | 2.46 | 0.66 |
| BPN-0037644 | 20.01 | 0.66 | 1.74 | 4.28 |
| BPN-0037655 | 24.89 | 0.19 | 0.47 | 0.23 |
| BPN-0036118 | 144.40 | 0.62 | 11.62 | 98.36 |
| BPN-0036610 | NC/I | 0.05 | 0.50 | 0.96 |
| BPN-0036611 | 4.72 | 0.62 | 1.02 | NC/I |
| BPN-0037549 | NC/I | 0.88 | 4.79 | 1.01 |
| BPN-0034927 | NC/I | 1.60 | 5.96 | 22.56 |
| BPN-0036015 | 39.13 | 2.09 | 67.30 | NC/I |
| BPN-0036195 | 178.00 | 28.30 | 438.70 | NC/I |
| BPN-0035940 | 70.77 | 1.96 | NC/I | NC/I |
| ROCKi | NC/I | 0.18 | 258.50 | NC/I |
| BPN-0035923 | 229.20 | 76.84 | NC/I | NC/I |
| S6K1i | 0.65 | NC/I | NC/I | NC/I |
| BPN-0036451 | NC/I | 2.88 | NC/I | NC/I |
| BPN-0036161 | NC/I | 56.88 | NC/I | NC/I |
| BPN-0036016 | 68.02 | NC/I | NC/I | NC/I |
| BPN-0036014 | NC/I | NC/I | NC/I | NC/I |
| BPN-0036402 | NC/I | NC/I | NC/I | NC/I |

NC/I = Not calculated/inactive

**Extended Data Table 7:** Phosphoproteomics Pathway Categories r.

| 2hr |  |  |  |  |  |  |  |  |  |  |  |  |  |  |  |
| --- | --- | --- | --- | --- | --- | --- | --- | --- | --- | --- | --- | --- | --- | --- | --- |
| Category | S6K1i_Avg(-LogP) | ROCKi_Avg(-LogP) | Combo_Avg(-LogP) | RO48_Avg(-LogP) | TMP316_Avg(-LogP) | S6K1i_AvgEnrichX_In(N+1) | ROCKi_AvgEnrichX_In(N+1) | Combo_AvgEnrichX_In(N+1) | RO48_AvgEnrichX_In(N+1) | TMP316_AvgEnrichX_In(N+1) | S6K1i_N | ROCKi_N | Combo_N | RO48_N | TMP316_N |
| PI3K/PTEN signaling pathways (30) | 3.45 | 3.31 | 3.02 | 3.65 | 3.90 | 5.43 | 7.11 | 6.60 | 9.32 | 11.11 | 6.00 | 6.00 | 11.00 | 13.00 | 22.00 |
| Rho GTPase signaling pathways (14) | 8.55 | 7.61 | 10.03 | 8.77 | 10.01 | 8.84 | 8.83 | 9.18 | 8.06 | 8.78 | 12.00 | 12.00 | 13.00 | 11.00 | 11.00 |
| Rac/Cdc42 signaling pathways (9) | 5.43 | 2.93 | 4.54 | 3.54 | 4.94 | 5.24 | 4.75 | 5.08 | 5.67 | 5.98 | 4.00 | 5.00 | 4.00 | 5.00 | 4.00 |
| Actin cytoskeleton pathways (23) | 5.00 | 4.41 | 5.46 | 6.02 | 6.28 | 7.87 | 6.40 | 6.42 | 8.43 | 9.91 | 19.00 | 15.00 | 16.00 | 21.00 | 21.00 |
| Microtubule cytoskeleton pathways (32) | 7.51 | 8.55 | 8.17 | 6.86 | 6.70 | 15.06 | 14.35 | 14.36 | 13.36 | 13.11 | 24.00 | 24.00 | 25.00 | 24.00 | 24.00 |
| Growth cone/presynaptic pathways (23) | 8.61 | 8.16 | 7.78 | 6.90 | 8.10 | 13.23 | 11.24 | 12.34 | 12.62 | 11.15 | 15.00 | 16.00 | 17.00 | 19.00 | 17.00 |
| Neurite outgrowth pathways (46) | 6.65 | 6.84 | 6.53 | 6.75 | 6.80 | 10.49 | 10.57 | 9.00 | 11.68 | 10.68 | 27.00 | 29.00 | 25.00 | 31.00 | 32.00 |
| Axon guidance pathways (8) | 4.77 | 5.74 | 5.79 | 4.11 | 8.35 | 8.18 | 6.60 | 9.53 | 5.41 | 5.38 | 4.00 | 5.00 | 5.00 | 5.00 | 3.00 |
| Neurotrophin signaling pathways (14) | 4.01 | 5.19 | 4.14 | 5.27 | 4.39 | 7.83 | 7.74 | 9.07 | 7.88 | 11.97 | 7.00 | 5.00 | 6.00 | 8.00 | 12.00 |
| Autophagy pathways (18) | 3.50 | 3.55 | 4.56 | 3.23 | 4.62 | 6.79 | 7.55 | 8.42 | 6.07 | 7.20 | 12.00 | 12.00 | 14.00 | 12.00 | 12.00 |
| Translation control pathways (18) | 3.22 | 3.97 | 4.40 | 3.44 | 4.13 | 5.90 | 7.08 | 7.09 | 6.39 | 6.59 | 8.00 | 9.00 | 11.00 | 10.00 | 10.00 |
| Vesicle trafficking pathways (66) | 4.01 | 4.28 | 3.93 | 3.99 | 4.15 | 14.48 | 15.95 | 12.90 | 12.01 | 15.41 | 38.00 | 43.00 | 33.00 | 36.00 | 51.00 |
| Endocytosis pathways (11) | 3.15 | 3.59 | 3.31 | 3.14 | 3.78 | 6.07 | 5.51 | 4.41 | 4.93 | 7.46 | 8.00 | 6.00 | 4.00 | 6.00 | 9.00 |
| Golgi/secretory pathways (20) | 3.40 | 3.62 | 3.67 | 2.92 | 3.63 | 6.85 | 7.65 | 8.94 | 6.89 | 7.46 | 8.00 | 9.00 | 13.00 | 8.00 | 10.00 |
| Focal adhesion/ECM pathways (16) | 2.63 | 2.39 | 2.63 | 2.68 | 3.03 | 7.09 | 3.25 | 3.26 | 4.38 | 7.05 | 3.00 | 3.00 | 3.00 | 7.00 | 7.00 |
| Cell polarity/Hippo pathways (27) | 4.28 | 4.78 | 5.41 | 5.13 | 4.88 | 9.75 | 9.49 | 9.71 | 9.58 | 12.25 | 14.00 | 14.00 | 12.00 | 12.00 | 16.00 |

| 24hr |  |  |  |  |  |  |  |  |  |  |  |  |  |  |  |
| --- | --- | --- | --- | --- | --- | --- | --- | --- | --- | --- | --- | --- | --- | --- | --- |
| Category | S6K1i_Avg(-LogP) | ROCKi_Avg(-LogP) | Combo_Avg(-LogP) | RO48_Avg(-LogP) | TMP316_Avg(-LogP) | S6K1i_AvgEnrichX_In(N+1) | ROCKi_AvgEnrichX_In(N+1) | Combo_AvgEnrichX_In(N+1) | RO48_AvgEnrichX_In(N+1) | TMP316_AvgEnrichX_In(N+1) | S6K1i_N | ROCKi_N | Combo_N | RO48_N | TMP316_N |
| PI3K/PTEN signaling pathways (30) | 3.50 | 2.88 | 3.23 | 3.78 | 3.77 | 9.05 | 7.00 | 8.34 | 12.05 | 12.66 | 17.00 | 11.00 | 18.00 | 20.00 | 18.00 |
| Rho GTPase signaling pathways (14) | 7.89 | 9.31 | 8.79 | 9.50 | 8.61 | 8.34 | 8.59 | 8.12 | 8.80 | 8.98 | 12.00 | 13.00 | 13.00 | 13.00 | 13.00 |
| Rac/Cdc42 signaling pathways (9) | 5.22 | 4.28 | 3.89 | 4.57 | 3.48 | 5.04 | 5.39 | 6.53 | 5.16 | 5.82 | 4.00 | 5.00 | 6.00 | 4.00 | 4.00 |
| Actin cytoskeleton pathways (23) | 4.90 | 5.27 | 5.12 | 5.23 | 4.58 | 6.42 | 4.81 | 6.02 | 5.66 | 7.23 | 15.00 | 10.00 | 16.00 | 14.00 | 20.00 |
| Microtubule cytoskeleton pathways (32) | 7.58 | 7.70 | 8.03 | 7.60 | 6.03 | 14.02 | 12.74 | 14.87 | 13.35 | 11.74 | 24.00 | 23.00 | 27.00 | 23.00 | 23.00 |
| Growth cone/presynaptic pathways (23) | 6.92 | 7.54 | 6.72 | 7.82 | 7.10 | 10.86 | 11.86 | 12.57 | 12.29 | 13.83 | 17.00 | 18.00 | 20.00 | 18.00 | 20.00 |
| Neurite outgrowth pathways (46) | 7.13 | 7.42 | 6.96 | 6.53 | 6.56 | 12.24 | 11.77 | 12.54 | 12.47 | 11.16 | 30.00 | 32.00 | 38.00 | 36.00 | 32.00 |
| Axon guidance pathways (8) | 3.99 | 4.74 | 4.51 | 4.93 | 5.46 | 8.46 | 8.32 | 8.51 | 5.80 | 6.17 | 6.00 | 7.00 | 7.00 | 5.00 | 5.00 |
| Neurotrophin signaling pathways (14) | 4.65 | 4.14 | 4.00 | 3.95 | 4.11 | 8.50 | 7.94 | 8.54 | 7.85 | 9.05 | 8.00 | 8.00 | 8.00 | 8.00 | 8.00 |
| Autophagy pathways (18) | 3.22 | 2.78 | 3.84 | 4.13 | 3.21 | 6.96 | 7.19 | 6.54 | 6.68 | 6.57 | 14.00 | 10.00 | 11.00 | 12.00 | 12.00 |
| Translation control pathways (18) | 3.69 | 3.69 | 4.02 | 3.74 | 4.16 | 9.60 | 8.60 | 5.04 | 7.29 | 8.87 | 9.00 | 11.00 | 7.00 | 10.00 | 11.00 |
| Vesicle trafficking pathways (66) | 3.84 | 3.99 | 3.96 | 4.10 | 4.11 | 15.00 | 13.53 | 13.40 | 14.20 | 14.78 | 44.00 | 40.00 | 41.00 | 43.00 | 42.00 |
| Endocytosis pathways (11) | 3.01 | 2.94 | 2.87 | 3.15 | 4.45 | 3.95 | 4.71 | 4.19 | 4.62 | 4.79 | 5.00 | 7.00 | 6.00 | 6.00 | 3.00 |
| Golgi/secretory pathways (20) | 2.69 | 3.06 | 3.18 | 4.35 | 3.51 | 10.05 | 9.96 | 6.60 | 7.42 | 5.80 | 9.00 | 11.00 | 10.00 | 9.00 | 7.00 |
| Focal adhesion/ECM pathways (16) | 2.86 | 2.65 | 2.93 | 2.92 | 3.46 | 3.45 | 6.45 | 6.90 | 6.79 | 6.22 | 3.00 | 6.00 | 9.00 | 7.00 | 9.00 |
| Cell polarity/Hippo pathways (27) | 5.26 | 4.76 | 4.56 | 5.32 | 5.46 | 8.53 | 7.97 | 11.69 | 10.66 | 10.63 | 9.00 | 10.00 | 15.00 | 11.00 | 12.00 |

**Extended Data Table 8:** Phosphosites with known functional relevance to neurite/axon growth.

| Gene | Site | Functions | Refs (PMIDs) |
| --- | --- | --- | --- |
| AKT1/2/3 | S473/S474/S472 | PI3K/PTEN signaling. Increased phosphorylation indicates PI3K activation and/or PTEN inhibition. | 15718470, 8978681, 28626016 |
| BASP1 | S40, S170 | Neuroprotection; actin polymerization; Ca<sup>2+</sup>-triggered mitochondrial dynamics in growing neurites. | 29229832, 33015061 |
| CRMP1 | T509, S518, S522, S523 | Pro-growth cytoskeletal activity modulation; MT dynamics at growth cone. | 18003833, 15676027 |
| DPYSL2 | S514, S518, S522 | Tubulin binding and axon elongation; tubulin binding and axonal guidance signaling modulation. | 15652488, 15676027 |
| GAP43 | S142 | JNK1-dependent; axonal growth; cytoskeletal remodeling. | 35347634 |
| GAP43 | S96 | Axon growth; axon regeneration; actin cytoskeletal remodeling. | 30240740 |
| MARCKS | S163 | PKC-phosphorylation mediated membrane dissociation; actin and PIP2 regulation; cell motility; adhesion; exocytosis; axon growth modulation. | 8557118 |
| PAK1 | T212 | Phosphorylated by p35/Cdk5 in postmitotic neurons. Rac1-dependent. Modulates microtubule remodeling at axonal growth cones during neurite outgrowth. | 11604394, 12176334 |
| PAK2 | S141 | Phosphorylation triggered by Rac1/Cdc42 activity. Modulates cytoskeletal remodeling at growth cones. | 16204230 |
| RPS6 | S236 | Selective mRNA translation; synaptic plasticity. Phosphorylated by S6K1/2 and RSK1/2. | 16166381, 28626016 |
| RPS6 | S240 | Selective mRNA translation; synaptic plasticity, neurite outgrowth. Phosphorylated by S6K1/2. | 16166381, 28626016 |
| STMN1 | S16 | Stathmin inactivation; MT stabilization; Ca<sup>2+</sup> signaling; neuronal polarization; axon specification. | 19359244 |
| STMN1 | S25 | Stathmin inactivation. Growth factor signaling; axon elongation. Erk1/2 substrate. | 8125092, 8325880 |
| STMN1 | S38 | Stathmin inactivation; axonal growth; growth cone dynamics. | 8125092 |
| STMN1 | S63 | Stathmin inactivation; MT stability; cAMP signaling. | 19359244 |
| STMN2 | S95 (S50) | MT stability; neuronal growth. | 14598370, 16838365 |

## References

1. Ahuja, C. S., et al. Traumatic spinal cord injury. Nat. Rev. Dis. Primers 3, 17018-(2017).

2. Park, K. K. et al. Promoting axon regeneration in the adult CNS by modulation of the PTEN/mTOR pathway. Science 322, 963–6 (2008).

3. Liu, K. et al. PTEN deletion enhances the regenerative ability of adult corticospinal neurons. Nat. Neurosci. 13, 1075–1081 (2010).

4. Sun, F. et al. Sustained axon regeneration induced by co-deletion of PTEN and SOCS3. Nature 480, 372–375 (2011).

5. Scheuren, P. S. & Kramer, J. L. K. Next-gen spinal cord injury clinical trials: lessons learned and opportunities for future success. EBioMedicine 109, (2024).

6. Kitano, H. Biological robustness. Nat. Rev. Genet. 5, 826–837 (2004).

7. Hopkins, A. L. Network pharmacology: the next paradigm in drug discovery. Nature Chemical Biology 2008 4:11 4, 682–690 (2008).

8. Ryszkiewicz, P., Malinowska, B. & Schlicker, E. Polypharmacology: new drugs in 2023-2024. Pharmacol. Rep. 77, 543–560 (2025).

9. Al-Ali, H. et al. Rational Polypharmacology: Systematically Identifying and Engaging Multiple Drug Targets to Promote Axon Growth. ACS Chem. Biol. 10, 1939–1951 (2015).

10. Gautam, P., Jaiswal, A., Aittokallio, T., Al-Ali, H. & Wennerberg, K. Phenotypic Screening Combined with Machine Learning for Efficient Identification of Breast Cancer-Selective Therapeutic Targets. Cell Chem. Biol. 26, 970–979.e4 (2019).

11. Patel, A. K. et al. Inhibition of GCK-IV kinases dissociates cell death and axon regeneration in CNS neurons. Proceedings of the National Academy of Sciences 117, 33597–33607 (2020).

12. Zheng, B. & Tuszynski, M. H. Regulation of axonal regeneration after mammalian spinal cord injury. Nat. Rev. Mol. Cell Biol. 24, 396–413 (2023).

13. He, Z. & Jin, Y. Intrinsic Control of Axon Regeneration. Neuron 90, 437–451 (2016).

14. Mah, K. M. et al. Compounds co-targeting kinases in axon regulatory pathways promote regeneration and behavioral recovery after spinal cord injury in mice. Exp. Neurol. 355, 114117 (2022).

15. Al-Ali, H. et al. Scaffold Ranking and Positional Scanning Identify Novel Neurite Outgrowth Promoters with Nanomolar Potency. ACS Med. Chem. Lett. 9, 1057–1062 (2018).

16. Kwon, B. K. et al. Intrathecal pressure monitoring and cerebrospinal fluid drainage in acute spinal cord injury: a prospective randomized trial. J. Neurosurg. Spine 10, 181–193 (2009).

17. Wu, D. et al. The blood–brain barrier: structure, regulation, and drug delivery. Signal Transduct. Target. Ther. 8, 217- (2023).

18. Awada, B., et al. Phenotypic Screening with Primary and Human iPSC-derived Neurons. Assay Guidance Manual https://www.ncbi.nlm.nih.gov/books/NBK619698/ (2025).

19. Vasta, J. D. et al. Quantitative, Wide-Spectrum Kinase Profiling in Live Cells for Assessing the Effect of Cellular ATP on Target Engagement. Cell Chem. Biol. 25, 206–214.e11 (2018).

20. Fournier, A. E., Takizawa, B. T. & Strittmatter, S. M. Rho kinase inhibition enhances axonal regeneration in the injured CNS. J. Neurosci. 23, 1416–23 (2003).

21. Al-Ali, H. et al. The mTOR Substrate S6 Kinase 1 (S6K1) Is a Negative Regulator of Axon Regeneration and a Potential Drug Target for Central Nervous System Injury. The Journal of Neuroscience 37, 7079–7095 (2017).

22. Anastassiadis, T., Deacon, S. W., Devarajan, K., Ma, H. & Peterson, J. R. Comprehensive assay of kinase catalytic activity reveals features of kinase inhibitor selectivity. Nat. Biotechnol. 29, 1039–45 (2011).

23. Manning, G., Whyte, D. B., Martinez, R., Hunter, T. & Sudarsanam, S. The protein kinase complement of the human genome. Science 298, 1912–34 (2002).

24. Pearce, L. R. et al. Characterization of PF-4708671, a novel and highly specific inhibitor of p70 ribosomal S6 kinase (S6K1). Biochem. J. 431, 245–55 (2010).

25. Tamura, M. et al. Development of specific Rho-kinase inhibitors and their clinical application. Biochim. Biophys. Acta Proteins Proteom. 1754, 245–252 (2005).

26. Kunkel, J., Luo, X. & Capaldi, A. P. Integrated TORC1 and PKA signaling control the temporal activation of glucose-induced gene expression in yeast. Nature Communications 2019 10:1 10, 3558- (2019).

27. Levin, D. E. Cell Wall Integrity Signaling in Saccharomyces cerevisiae. Microbiology and Molecular Biology Reviews 69, 262 (2005).

28. Kamada, Y. et al. Tor2 Directly Phosphorylates the AGC Kinase Ypk2 To Regulate Actin Polarization. Mol. Cell. Biol. 25, 7239 (2005).

29. Johnson, J. L. et al. An atlas of substrate specificities for the human serine/threonine kinome. Nature 613, 759–766 (2023).

30. Um, S. H. et al. Absence of S6K1 protects against age- and diet-induced obesity while enhancing insulin sensitivity. Nature 2004 431:7005 431, 200–205 (2004).

31. Sarbassov, D. D., Guertin, D. A., Ali, S. M. & Sabatini, D. M. Phosphorylation and regulation of Akt/PKB by the rictor-mTOR complex. Science (1979). 307, 1098–1101 (2005).

32. Li, Z. et al. Regulation of PTEN by Rho small GTPases. Nature Cell Biology 2005 7:4 7, 399–404 (2005).

33. Manser, E., Leung, T., Salihuddin, H., Zhao, Z. S. & Lim, L. A brain serine/threonine protein kinase activated by Cdc42 and Rac1. Nature 1994 367:6458 367, 40–46 (1994).

34. Hartwig, J. H. et al. MARCKS is an actin filament crosslinking protein regulated by protein kinase C and calcium–calmodulin. Nature 1992 356:6370 356, 618–622 (1992).

35. Dunham, K. A., Siriphorn, A., Chompoopong, S. & Floyd, C. L. Characterization of a graded cervical hemicontusion spinal cord injury model in adult male rats. J. Neurotrauma 27, 2091–2106 (2010).

36. Spinal Cord Injury Statistical Center in collaboration with the Model Systems Knowledge Translation Center, N. Spinal Cord Injury Facts and Figures at a Glance 2020 SCI Data Sheet. www.msktc.org/sci/model-system-centers. (2020).

37. Lehmann, M. et al. Inactivation of Rho Signaling Pathway Promotes CNS Axon Regeneration. Journal of Neuroscience 19, 7537–7547 (1999).

38. Sivasankaran, R. et al. PKC mediates inhibitory effects of myelin and chondroitin sulfate proteoglycans on axonal regeneration. Nat. Neurosci. 7, 261–268 (2004).

39. Wang, X., Hu, J., She, Y., Smith, G. M. & Xu, X. M. Cortical PKC inhibition promotes axonal regeneration of the corticospinal tract and forelimb functional recovery after cervical dorsal spinal hemisection in adult rats. Cerebral Cortex 24, 3069–3079 (2014).

40. Wei, D. et al. Inhibiting cortical protein kinase A in spinal cord injured rats enhances efficacy of rehabilitative training. Exp. Neurol. 283, 365–374 (2016).

41. Hofmann, F., Feil, R., Kleppisch, T. & Schlossmann, J. Function of cGMP-dependent protein kinases as revealed by gene deletion. Physiol. Rev. 86, 1–23 (2006).

42. Buchser, W. J., Slepak, T. I., Gutierrez-Arenas, O., Bixby, J. L. & Lemmon, V. P. Kinase/phosphatase overexpression reveals pathways regulating hippocampal neuron morphology. Mol. Syst. Biol. 6, 391 (2010).

43. Anighoro, A., Bajorath, J. & Rastelli, G. Polypharmacology: Challenges and opportunities in drug discovery. J. Med. Chem. 57, 7874–7887 (2014).

44. Jia, J. et al. Mechanisms of drug combinations: interaction and network perspectives. Nature Reviews Drug Discovery 2009 8:2 8, 111–128 (2009).

45. Hilton, B. J. et al. An active vesicle priming machinery suppresses axon regeneration upon adult CNS injury. Neuron 110, 51–69.e7 (2022).

46. He, M. et al. Autophagy induction stabilizes microtubules and promotes axon regeneration after spinal cord injury. Proc. Natl. Acad. Sci. U. S. A. 113, 11324–11329 (2016).

47. Lowell, J. A. et al. Phenotypic Screening Following Transcriptomic Deconvolution to Identify Transcription Factors Mediating Axon Growth Induced by a Kinase Inhibitor. SLAS Discovery 26, 1337–1354 (2021).

48. Sadybekov, A. V. & Katritch, V. Computational approaches streamlining drug discovery. Nature 616, 673–685 (2023).

49. Gustafsson, J. R., Katsioudi, G., Issazadeh-Navikas, S. & Kornum, B. R. Neurobasal media facilitates increased specificity of siRNA-mediated knockdown in primary cerebellar cultures. J. Neurosci. Methods 274, 116–124 (2016).

50. Nagendran, T., Poole, V., Harris, J. & Taylor, A. M. Use of Pre-Assembled Plastic Microfluidic Chips for Compartmentalizing Primary Murine Neurons. J. Vis. Exp. 2018, (2018).

51. Wu, C., Schulte, J., Sepp, K. J., Littleton, J. T. & Hong, P. Automatic robust neurite detection and morphological analysis of neuronal cell cultures in high-content screening. Neuroinformatics 8, 83–100 (2010).

52. Metz, J. T. et al. Navigating the kinome. Nat. Chem. Biol. 7, 200–2 (2011).

53. Passaro, S. et al. Boltz-2: Towards Accurate and Efficient Binding Affinity Prediction. *bioRxiv* 10.1101/2025.06.14.659707 (2025) doi:10.1101/2025.06.14.659707.

54. Elwardany, O. S., et al. Lumbar intrathecal catheterization in rats targeting the cerebral cortex: a drug delivery method and validation. *bioRxiv* 2026.06.01.727192 (2026) doi:10.64898/2026.06.01.727192.

55. Torres-Espín, A. et al. Promoting FAIR Data Through Community-driven Agile Design: the Open Data Commons for Spinal Cord Injury (odc-sci.org). Neuroinformatics 20, 203– 219 (2022).

56. Zörner, B. et al. Profiling locomotor recovery: comprehensive quantification of impairments after CNS damage in rodents. Nat. Methods 7, 701–708 (2010).

57. Christ, M., Braun, N., Neuffer, J. & Kempa-Liehr, A. W. Time Series FeatuRe Extraction on basis of Scalable Hypothesis tests (tsfresh – A Python package). Neurocomputing 307, 72–77 (2018).

58. Eid, S., Turk, S., Volkamer, A., Rippmann, F. & Fulle, S. KinMap: a web-based tool for interactive navigation through human kinome data. BMC Bioinformatics 2017 18:1 18, 16- (2017).

59. Graczyk, P. P. Gini coefficient: A new way to express selectivity of kinase inhibitors against a family of kinases. J. Med. Chem. 50, 5773–5779 (2007).

