## Supplementary Materials for "AGC kinase homology requires and enables co-targeting for CNS regeneration"

### Supplementary Methods

#### Chemicals and small molecule kinase inhibitors

RO48 and Blueprint Neurotherapeutics Network (BPN) program analogs were synthesized by Curia (Albany, NY, USA) and assigned BPN identifiers, e.g., BPN-0037316 (TMP-316) and BPN-0034858 (RO48). Earlier analogs were synthesized by Sanford Burnham Prebys Medical Discovery Institute (SBI, Lake Nona, FL, USA) and given SBI identifiers. The purity of TMP-316 used in animal studies was >99% by ultra-high-performance liquid chromatography. The clinical formulation of TMP-316 (20 mM) was prepared in 10% w/v 2-hydroxypropyl-β-cyclodextrin (Sigma-Aldrich, 332607), 6.125 mM $NaH_2PO_4$ (Sigma-Aldrich, >99%, S0876), 18.875 mM $Na_2HPO_4$ (Sigma-Aldrich, >99%, S0751), 48 mM NaCl (Fisher Scientific, ACS grade, >99%, AA1231436) in HPLC-grade water (Fisher Scientific, W5-4), pH 5. This formulation was developed by the BPN program (NIH/NINDS UH3NS124630) and was used in all in vivo studies. Formulated solutions were filtered through a 0.2 µm polytetrafluoroethylene (PTFE) filter (Millipore, SLFG025LS) immediately before use. Chemical syntheses of program compounds are reported in US Patent No. 11,926,613 B2. Detailed structure-activity relationships will be described elsewhere. Neurite outgrowth (NTL) data for all compounds used in this study are provided in Supplementary Tables 2 and 3. Rho Kinase Inhibitor IV (ROCKi) was purchased from Santa Cruz Biotechnology (SC-222254) and PF-4708671 (S6K1i) was purchased from MilliporeSigma (559273). All stock solutions for in vitro testing were prepared in dimethyl sulfoxide (DMSO; MilliporeSigma, D2650).

**Neuronal cultures and neurite outgrowth assays**

Rat primary hippocampal neurons and human iPSC-derived glutamatergic neurons were cultured and utilized in phenotypic assays as detailed in Awada et al.[1] Briefly, primary hippocampal neurons were isolated from E18 Sprague-Dawley rat embryos (pooled males and females) by trypsin/DNase I dissociation of dissected hippocampi, followed by mechanical trituration, and seeded in NbActiv4 (BrainBits, NB4) at assay-specific densities. Human iPSC-derived glutamatergic neurons (iGLUTs; ioGlutamatergic Neurons, bit.bio, io1001) were revived in Neurobasal medium (ThermoFisher, 21103049) supplemented with Glutamax (1x, ThermoFisher, 35050061) and 2-mercaptoethanol (25 µM, ThermoFisher, 31350010). iGLUTs were resuspended and seeded in media further supplemented with B27 (1x, ThermoFisher, 17504044), NT3 (10 ng/mL, R&D, 267-N3-005), BDNF (5 ng/mL, R&D, 248-BDB-005), and doxycycline (1 µg/mL, Sigma, D9891).

Rat primary hippocampal neurons or human iGLUTs were each seeded at densities of 2,000 or 4,000 cells/well, respectively, onto 96-well plates previously coated for 24 h with 0.5 mg/mL PDL (poly-D-lysine, Sigma-Aldrich, P6407) in HBSS (Hanks' Balanced Salt Solution, Sigma-Aldrich, H9394-24X500ML) with 20 mM HEPES (Gibco, 15630-080). Cultures were incubated at 37 °C with 5% $CO_2$ for 24 h (rat primary hippocampal neurons) or 72 h (iGLUTs), then treated with compounds for 48 h prior to fixation and immunostaining. For screening and SAR campaigns, RO48 was included on each plate as a benchmark for inter-run normalization, with three technical replicates performed per compound per run, with a minimum of two runs for each compound.

Neuronal cultures were fixed, permeabilized, and blocked in 4% PFA/PBS (20 min, 37°C), 0.3% Triton X-100/PBS (30 min), and 0.2% fish gelatin with 0.03% Triton X-100 in PBS (1 h), respectively, followed by overnight incubation at 4°C with anti-βIII-tubulin antibody (Sigma-Aldrich, T2200, RRID:AB_262133; 1:1,000) in blocking buffer. Plates were washed with PBS, and Alexa Fluor 488 goat anti-rabbit secondary antibody (Invitrogen, A11034, RRID:AB_2576217; 1:1,000) and Hoechst 33342 (Invitrogen, H3570; 10 μg/mL) in blocking buffer were applied for 1 h at room temperature. Plates were washed with PBS and imaged on the Opera Phenix High Content Screening (HCS) system, and Neurite total length (NTL) was quantified using Harmony software with automated segmentation of Hoechst-positive nuclei and βIII-tubulin-positive processes.

NTL values, computed as the mean per well, were normalized to DMSO vehicle controls within each plate and expressed as %NTL. The drug effect score (DES), defined as the area under the dose-response curve above baseline, was normalized to the RO48 DES within the same assay run to yield %DES[1].

To correct for inter-run variability in program-wide analyses, a benchmarked %NTL was calculated relative to the RO48 reference compound within each run. The maximum response of RO48 was set to 100%, and the baseline vehicle response to 0%, and benchmarked %NTL was calculated as follows:

$$\text{Benchmarked \%NTL} = 100 \times \frac{\%\text{NTL} - 100}{\text{Max}(\%\text{NTL}_{\text{RO48}}) - 100}$$

**Neuronal viability assay**

Primary hippocampal neurons were isolated from E18 Sprague-Dawley rat embryos and cultured as described above. Neurons were seeded at 7,500 cells/well in poly-D-lysine-coated 96-well plates and allowed to recover for 1 h prior to compound addition. Compounds were added in dose-response format and cells were incubated for 48 h. Each run included vehicle controls and a positive cell-death control (FLT-3 inhibitor). Two technical replicates were performed per compound per run. Cell viability was assessed using the CellTiter-Glo Luminescent Cell Viability Assay (Promega). At the assay endpoint, medium was aspirated leaving approximately 40 µL per well, 40 µL CellTiter-Glo reagent was added, and plates were shaken for 2 min at 500 rpm to induce cell lysis. Following a 15 min incubation at room temperature, luminescence was measured on a GloMax Discover microplate reader (Promega). Viability values were normalized to vehicle-treated controls and expressed as percent viability.

**Validation of siRNA-mediated gene silencing**

Neurons were seeded at a density of 250,000 cells/well in 48-well plates and allowed to adhere overnight before being treated with 1 µM of Scramble, α-PKCγ, or α-PKX siRNA for 48 h. Culture medium was then removed, neurons were washed with PBS, and total RNA was extracted using the Arcturus® PicoPure® RNA Isolation Kit (Applied Biosystems). RNA was quantified and used as a template for cDNA synthesis using the Omniscript RT kit (Qiagen), random primers (Promega), and RNAse inhibitor (New England Biolabs). PKCγ and PKX mRNA levels were evaluated via reverse transcription quantitative PCR (RT-qPCR) using PowerUp™ SYBR™ Green Master Mix (Applied Biosystems) and the QuantStudio™ 3 Real-Time PCR System (both Applied Biosystems). All reactions were performed in a final volume of 10 µL with cycling conditions of UNG activation at 50°C for 2 min, initial denaturation at 95°C for 2 min, 40 cycles of 15 s denaturation at 95°C and 1 min extension at 60°C, with a final melt curve analysis (95°C for 15 s, 60°C for 1 min, 95°C for 15 s). Fold-change for each gene was then calculated according to the Livak ($2^{-\Delta\Delta Ct}$) method[2]. Data were expressed as percentage of the scramble siRNA control, with GAPDH used as a reference gene. Primer sequences for GAPDH and PKX are listed below. For PKCγ, the commercially available PrimePCR AssayPrkcg (BIO-RAD, qRnoCID0002483) was used.

| Primer Name | Sequence (5'→3') |
|---|---|
| GAPDH Forward | GGGTGTGAACCACGAGAAAT |
| GAPDH Reverse | ACTGTGGTCATGAGCCCTTC |
| PKX Forward | CCACACCACACTGATGACCA |
| PKX Reverse | TGGGCAGGTTTCCAGTTGTT |

**Microfluidic axotomy assay**

Pre-assembled two-compartment microfluidic devices (XonaChip® 450 μm barrier, Xona Microfluidics, XC450) were used to physically isolate neuronal soma from axons, as previously described[3]. Briefly, channels were sequentially pre-coated with 0.5 mg/mL PDL for 1 h at 37 °C according to the manufacturer's instructions and equilibrated with NbActiv4 medium. E18 hippocampal neurons were seeded at a density of $1.2 \times 10^5$ cells per device by loading 5 μL cell suspension into both channels of the somatic compartment. Cell cultures were maintained at 37 °C with 5% $CO_2$, and the medium was changed every 2 days.

At 8 DIV, axotomy was performed by complete vacuum aspiration (1-2 min) of the axonal compartment, with successful transection verified visually under a microscope. Medium in the somatic compartment, the axonal compartment, or both was replaced with NbActiv4 containing RO48 [2 μM] or DMSO vehicle. Following a 48-h incubation, neurons were fixed in 4% PFA (30 min), permeabilized in 0.25% Triton X-100/PBS (15 min), and blocked in 10% normal goat serum (Sigma-Aldrich; 15 min). Immunostaining was performed overnight at 4°C with anti-βIII-tubulin in 1% goat serum, followed by incubation with Alexa Fluor 488-conjugated goat anti-rabbit secondary antibody (1 h, room temperature).

Devices were imaged on a Dragonfly spinning disk confocal microscope (Andor/Oxford Instruments) using a 20X objective. Tile scans of the axonal compartments were acquired, and Z-stacks were collected at 1 μm z-intervals; maximum intensity projection images were generated for analysis**.** Axon regrowth was quantified using an automated tracing MATLAB (R2025b) script modified from Wu et al.[4] Values from each run were converted to robust z-scores as previously described.

**Bliss independence synergy analysis**

The combined effect of ROCKi and S6K1i on neurite outgrowth was evaluated in the primary neuron NTL assay using a 5×5 concentration matrix (0, 1.25, 2.5, 5, and 10 μM each), with 3 technical replicates per condition. For each condition, effect size was calculated as %NTL – 100, then normalized to a 0–1 scale by dividing by the effect size of the highest combination. Synergy was quantified using the Bliss independence model[5]: $E_{A,B} = E_A + E_B - E_A \times E_B$, where $E_A$ and $E_B$ are the normalized single-agent effects from the corresponding 0 μM arms of the matrix. The Bliss synergy score was $E_{Combo} - E_{A,B}$. Positive scores indicate synergy; negative scores indicate antagonism.

**Random forest modeling**

Random forest regression was used to assess the cross-validated predictive contribution of each kinase to Benchmarked NTL. Per-observation target engagement (%TE) values for S6K1, ROCK2, PKX, and PKCγ were used as features; Benchmarked NTL was the response. All combinations of 1, 2, 3, and 4 kinases (15 total) were evaluated, along with three derived single-feature summaries (minimum, sum, and product of TE across the 4 kinases). For each combination, a random forest of 100 trees with a maximum depth of 10 was trained using MATLAB's (R2025b) TreeBagger. Performance was estimated by 5-fold cross-validation with a fixed random seed (rng = 42). Performance was quantified using the coefficient of determination ($R^2$) calculated on held-out test folds, with mean cross-validated $R^2$ used to compare feature sets.

**Multiple kinase domain sequence alignments**

Full-length human protein sequences were retrieved from UniProtKB (release 2026_01) for ROCK2 (O75116), PKCγ (PRKCG; P05129), S6K1 (RPS6KB1; P23443) and PKX (P51817). For each protein, the catalytic domain was excised according to the boundaries of the UniProt-annotated "Protein kinase" domain, giving residues 92–354 of ROCK2 (263 aa), 351–614 of PKCγ (264 aa), 91–352 of S6K1 (262 aa) and 49–303 of PKX (255 aa). The four kinase domain sequences were aligned using Clustal Omega v1.2.4 with default parameters.

**Lineage analysis of TMP-316's AGC kinase targets**

The evolutionary origins of ROCK2, S6K1, PKCγ, and PKX were traced across Saccharomyces cerevisiae, Caenorhabditis elegans, Drosophila melanogaster, Rattus norvegicus, and Homo sapiens, following the kinase classification framework of Manning et al.[6] Approximate divergence times are from TimeTree 5[7]. ROCK2 has no yeast ortholog; its closest invertebrate orthologs are let-502 (UniProtKB P92199; C. elegans) and Rok (UniProtKB Q9VXE3; Drosophila). The yeast gene PKC1 is included as the founding fungal member of the PKC clade but is phylogenetically placed within the PKN/PRK family and is not a conventional PKC ortholog. The closest invertebrate representatives of the PKC clade are tpa-1 (UniProtKB

P34722; C. elegans, novel PKC) and Pkc53E (UniProtKB P05130; Drosophila, conventional PKC); conventional PKC isoforms are a metazoan-specific expansion. S6K1 is the only target with a true yeast ortholog: ypk3, whose deletion abolishes ribosomal S6 phosphorylation and is functionally complemented by human S6K1. Its invertebrate orthologs are rsks-1 (C. elegans) and dS6K (Drosophila); rskn-1/2 (C. elegans) and dRSK (Drosophila) represent co-expanded RSK-related kinases in the same clade. PKX has no yeast ortholog; TPK1/2/3 are the closest yeast members (classical PKA catalytic subunits). Its invertebrate orthologs are F47F2.1 (NCBI Gene 180673; C. elegans) and DC2 (Drosophila); egl-4 (NCBI Gene 176991; C. elegans) and kin-1 (UniProtKB P21137; C. elegans) represent co-expanded PKG and PKA members, and foraging, DC0, and DC1 represent the equivalent Drosophila clade members.

**Computational docking**

Protein-ligand complex structures were co-folded using Boltz-2 (v2.2.0)[8]. Two structural templates were supplied per complex to guide co-folding: the target model obtained from the AlphaFold Protein Structure Database (AF-P23443-4-F1 for S6K1, AF-O75116-F1 for ROCK2, AF-P05129-2-F1 for PKCγ, AF-P51817-F1 for PKX), and a crystal structure of PKA catalytic subunit in complex with RO48 (PDB: 1SVG, 2.02 A resolution). Protein-ligand complexes were further optimized using a simulated annealing routine in OpenMM (v8.4.0). Structural representations of the protein-ligand complex were generated using PyMOL Molecular Graphics System (Schrödinger, LLC).

The structure of TMP-316 was provided as a SMILES string in the Boltz-2 input specification for co-folding with each of the four kinase targets (S6K1, ROCK2, PKCγ, and PKX). Predictions were performed using the following parameters: 3 recycling steps, 200 diffusion sampling steps, and 1 diffusion sample per target. Multiple sequence alignments were generated via the Boltz

MSA server. Affinity prediction with molecular weight correction was enabled. The top predicted pose was accepted per protein-ligand pair.

**Molecular dynamics**

Protein structures were prepared using PDBFixer at pH 7.0. The ligand was extracted from each Boltz-2 output using RDKit (v2025.09.4); bond orders were assigned from the SMILES representation via template matching, and hydrogens were added with 3D coordinate generation. The ligand was parameterized with the GAFF 2.11 force field via the OpenFF Toolkit and GAFFTemplateGenerator. The protein was described with the AMBER14 force field. Solvation was treated implicitly using the GBn2 generalized Born model. Systems were constructed using a nonperiodic cutoff scheme with a 2.0 nm nonbonded cutoff. Covalent bonds to hydrogen were constrained using the SHAKE algorithm.

Three classes of restraints were applied. First, harmonic positional restraints (1000 kJ/mol/$nm^2$) were applied to backbone atoms (N, Cα, C, O) of residues beyond 8 Å from the ligand, allowing local flexibility around the binding site while maintaining the overall protein fold. Second, soft harmonic positional restraints (10 kJ/mol/$nm^2$) were applied to ligand heavy atoms to prevent large-scale drift from the predicted binding pose. Third, distance restraints (1000 kJ/mol/$nm^2$, 2.5–3.5 Å bounds) were applied to enforce two conserved pharmacophoric interactions: a hydrogen bond between the ligand and the hinge region backbone amide, and a contact between the ligand and the sidechain amine of the catalytic lysine. The specific restrained residues corresponded to structurally equivalent positions across all four kinases.

Each system was subjected to an initial energy minimization (2000 steps), followed by a three-phase simulated annealing protocol: heating from 300 K to 500 K over 50,000 steps (100 ps), a high-temperature phase at 500 K for 100,000 steps (200 ps), and cooling from 500 K to 300 K over 150,000 steps (300 ps), totaling 0.6 ns. A final energy minimization (2000 steps) was performed on the annealed structure. The optimized coordinates were saved for downstream analysis. The structural models generated in this study have been deposited in ModelArchive[9] (https://modelarchive.org) with the accession code ma-dew4d.

**In vitro kinase assays**

TMP-316 and RO48 kinase inhibition profiling methods (KINOMEscan and Reaction Biology Corporation panels) are described in Main Methods. Profiling data for the inhibitors Y-27632 and Gö 6976 were obtained from the published dataset of Anastassiadis et al.[10], in which 178 compounds were profiled by Reaction Biology Corporation against a panel of 300 wild-type kinases at 500 nM in the presence of 10 μM ATP, in duplicate. Kinase activities were converted into percent inhibition and mapped onto the human kinome tree using KinMap[11]. Where a kinase was assayed under multiple conditions (for example, cyclin-dependent kinases tested with different cyclin partners, or splice variants), these entries were collapsed to a single kinome node represented by the maximum observed percent inhibition; kinase names were mapped to KinMap identifiers via their UniProt accessions, and assayed targets without a corresponding node on the kinome tree were not annotated.

**Phosphoproteomic analysis**

E18 rat hippocampal neurons were seeded at $4 \times 10^6$ cells per PDL-coated 60 mm culture dish in NbActiv4 and allowed to adhere overnight. Culture medium was replaced with fresh NbActiv4 containing TMP-316 [2 μM], RO48 [2 μM], ROCKi [10 μM], S6K1i [5 μM], or ROCKi+S6K1i. These concentrations were selected as those that maximized neurite outgrowth without causing cellular toxicity (>90% viability relative to vehicle) (Supplementary Fig. 1k). Cells were incubated for 2 or 24 h, washed with ice-cold PBS, scraped into ice-cold PBS supplemented with 1× ReadyShield protease and phosphatase inhibitor cocktail (Sigma-Aldrich), pelleted by centrifugation at 500 × g for 10 min at 4 °C, snap-frozen in liquid nitrogen, and stored at −80 °C. Each cell pellet was lysed in extraction buffer [7 M urea, 2 M thiourea, 0.4 M Tris pH 8, 20% acetonitrile, 10 mM tris(2-carboxyethyl)phosphine (TCEP), 40 mM chloroacetamide, and 1× HALT EDTA-free protease and phosphatase inhibitor (ThermoFisher Scientific)]. Samples were sonicated twice at 30% amplitude for 7 s with a Branson Digital Sonifier 250 (Branson Ultrasonics, Danbury, CT). Each sample was processed in a Barocycler NEP2320 (Pressure Biosciences, South Easton, MA) at 35 kpsi for 20 s and 0 kpsi for 10 s for 60 cycles at 37 °C. Lysates were transferred to 1.5 mL microfuge tubes and centrifuged at 15,000 × g for 10 min. An aliquot of each sample was taken for protein concentration determination by the Bradford assay. A 50 µg aliquot of each sample was transferred to a new 1.5 mL microfuge tube and brought to the same volume with extraction buffer. All samples were diluted fivefold with water, and then trypsin (Promega, Madison, WI) was added at a 1:40 ratio of enzyme-to-protein. Samples were incubated for 16 h at 37 °C. After incubation, each sample was acidified to 0.2% trifluoroacetic acid (TFA) to stop trypsin activity and brought to 500 µL with 0.1% TFA in water. Each sample was cleaned up with a 1 mL Waters Oasis HLB cartridge (Waters Corporation, Milford, MA), and the eluates were dried by vacuum centrifugation. Samples were enriched for phosphopeptides with the High-Select™ TiO2 phosphopeptide kit and the High-Select™ Fe-NTA phosphopeptide kit using the High-Select™ SMOAC (Sequential enrichment of Metal Oxide Affinity Chromatography) protocol per the manufacturer's instructions (ThermoFisher

Scientific, Rockford, IL). The resulting eluates were pooled and analyzed by data-independent acquisition on a Thermo Orbitrap Eclipse mass spectrometer.

Peptide identification and label-free quantification were performed in Proteome Discoverer 3.2.0.450 (Thermo Fisher Scientific). DIA spectra were searched with CHIMERYS (inferys_4.7.0_fragmentation prediction model) with match-between-runs against the rat reference proteome (UniProt UP000002494, released 2025-08-19). Trypsin was specified with full specificity and up to two missed cleavages; peptides were 7-30 residues with charge states 1-6. Precursor mass tolerance was 20 ppm, and fragment mass tolerance was 20 ppm. Carbamidomethylation of cysteine was set as a static modification; oxidation of methionine and phosphorylation of serine, threonine, and tyrosine were set as dynamic modifications, with a maximum of three dynamic modifications per peptide. Peptide-spectrum matches (PSMs) were filtered to a 1% false discovery rate (q-value). Peptides were likewise filtered to a 1% false discovery rate. Phosphosite assignment used a site probability threshold of 75%. Proteins were assembled under strict parsimony and filtered at 1% protein FDR; the reported protein set was restricted to master proteins with more than one unique peptide. Peptide quantification used MS2 fragment-ion apex intensities across all files, normalized to total peptide amount without imputation. Differential phosphorylation was assessed from pairwise abundance ratios computed on modified peptides relative to vehicle-treated controls at each time point (2 and 24 h), with significance evaluated by a background-based t-test. The experiment comprised six treatments (vehicle, TMP-316, RO48, ROCKi, S6K1i, and ROCKi+S6K1i) at 2 time points across three biological replicates derived from independent neuronal preparations. The mass spectrometry proteomics data have been deposited to the ProteomeXchange Consortium (http://proteomecentral.proteomexchange.org) via the PRIDE partner repository[12] with the dataset identifierPXD081808.

**Pathway analysis**

Differentially phosphorylated protein lists were generated for each treatment condition relative to vehicle by filtering the protein-level quantification (Supplementary Table 5) to entries carrying a phosphorylation modification; proteins without a phosphorylation annotation were excluded. Each treatment sample group was compared to its time-matched vehicle group, and for each comparison, the log2 fold change was calculated from the linear Abundance Ratio. Phosphorylated proteins with |log2 fold change| > 0.5 and an adjusted p-value < 0.1 were retained to maximize sensitivity, with more stringency imposed at the enrichment step.

The retained proteins were mapped to their gene symbols, and the resulting gene lists were submitted to Metascape (v3.5.20250701), an open-access web-based portal for gene-list enrichment analysis that integrates multiple biological knowledge bases. Gene identifiers were mapped to Rattus norvegicus Entrez gene IDs; all genes in the genome were used as the enrichment background, and protein-protein interaction analysis used the human interaction network. GO Biological Process terms with an enrichment factor > 1.5 and LogP < −2 were retained (Supplementary Tables 6 and 7).

Significant terms were partitioned into mutually exclusive biological categories defined a priori for this neurite-outgrowth model, with each term assigned to exactly one category. Terms not applicable to dissociated neurons (e.g. cancer, hepatic, renal) were excluded, and a subset of presynaptic and synaptic-vesicle terms were re-assigned as growth-cone and constitutive-trafficking processes (Extended Data Table 7). Terms with no biological relevance to the study (e.g., cancer, hepatic, renal, and cardiac annotations) were removed by keyword filtering. For each category-condition pair, N was the number of enriched terms, AvgEnrich the mean Metascape enrichment factor across those terms, and Avg(−LogP) the mean negative LogP across the same subset. A size-corrected composite score was calculated as AvgEnrich × ln(N + 1) to permit enrichment comparisons across categories with large size differences (Extended Data Table 7). Avg(−LogP) and the composite score are descriptive summaries used for cross-category comparison.

**TMP-316 in vivo pharmacokinetics**

The pharmacokinetic profile of TMP-316 following IT administration was assessed in male Sprague-Dawley rats (10–11 weeks old). In vivo work for this experiment was performed at AfaSci Research Laboratories (Redwood City, CA). Animals were assigned to six subgroups (n = 4 per subgroup) and administered a single IT injection at the L5–6 intervertebral space over a 4-min infusion period (40 µl per animal; ~1 mg/kg). Plasma, CSF, brain, and spinal cord samples were collected at 0.25, 0.5, 1, 1.5, 2, and 2.5 h post-dose. CSF was aspirated from the cisterna magna under 2.5% isoflurane, followed by terminal whole blood collection via cardiac puncture. Animals were subsequently euthanized, and brain and spinal cord tissues were collected, rinsed with cold saline, blotted dry, and frozen until analysis.

Bioanalytical quantification of TMP-316 in all matrices was performed at SRI International (Menlo Park, CA) by LC-MS/MS on an AB Sciex 5500 Q-Trap mass spectrometer, with chromatographic separation on a Phenomenex Synergi Polar RP column (2.0 × 100 mm, 4.0 µm), using propranolol as an internal standard. The lower limit of quantitation (LLOQ) was 1.00 ng/ml for plasma and CSF, and 5.00 ng/g for brain and spinal cord homogenate. Pharmacokinetic parameters were derived by noncompartmental analysis using Phoenix WinNonlin® (v8.5) with the sparse sampling feature.

#### In vivo target engagement

Adult male Sprague-Dawley rats (9 weeks old, ~300 g) were administered TMP-316 (20 mM formulation, ~1 mg/kg) or vehicle (formulation buffer) via lumbosacral IT delivery (30 µL at ~6 µL/min) as previously described by Elwardany et al[54]. Animals were randomly assigned to treatment groups and personnel were blinded to treatment allocation. At 1.5 h post-dosing (Tmax, Extended Data Table 3), animals were euthanized, CNS tissues were rapidly dissected, and 100 mg of cortical tissue samples were lysed in hot 8% SDS lysis buffer [Tris-HCl (0.2 M, pH 6.5), 32% v/v glycerol, 8% w/v SDS, 0.04% w/v bromophenol blue, with cOmplete Mini EDTA-free protease inhibitor cocktail (Roche)]. S6K1 activity was assessed by Western blot using antibodies against phospho-S6 (Ser240/244; Cell Signaling, 5364L, RRID:AB_10694233; 1:1,000) and total S6 (Cell Signaling, 2317S, RRID:AB_2238583; 1:500). Band intensities were quantified using AzureSpot Pro software. The pS6/S6 ratio was computed for each sample, normalized to vehicle-treated controls, and expressed as a % relative of vehicle control.

#### Analysis of kinematic data

Kinematic time-series data were exported from the MotoRater software as 46 pre-computed base parameters per trial. A custom preprocessing step was applied to each time series to remove leading and trailing segments of zero variance, retaining only the locomotor-active portion of each trial. Automated feature extraction was performed using the TSFresh library (v0.21.1)[13], which systematically computed hundreds of time-series descriptors per input signal spanning distributional, spectral, complexity, autocorrelation, and nonlinear features. Applied across all 46 base parameters, this yielded approximately 36,800 candidate features per trial. A two-stage relevance filtering procedure was then applied using TSFresh's hypothesis-testing framework with false discovery rate correction ($\alpha = 0.01$), evaluating each feature's ability to discriminate: (i) pre-injury baseline (0 DPI) from acute injury (7 DPI), and (ii) vehicle-treated from TMP-316-treated animals at post-injury time points (>7 DPI). Features surviving both filters were combined, yielding approximately 10,000 significant candidate features. From this pool, the top 500 features ranked by aggregated relevance p-value were selected, subject to a diversity constraint capping selection at 20 features per base parameter, yielding a final extracted feature set of 233 features derived from 25 of 46 base parameters. Features with absolute skewness exceeding 1.0 were normalized using the Yeo–Johnson transformation, after which all features were standardised to zero mean and unit variance (StandardScaler; scikit-learn v1.6.1)[scikit-learn_JMLR_2011_pedregosa]. Features were then filtered to retain only those with treatment-relevant signal in the post-injury period: a linear regression was fit to each animal's post-injury recovery trajectory (7–42 DPI) for each feature, with trial-level data averaged across runs prior to slope computation. Recovery slopes were compared across treatment groups using a one-way ANOVA, and features showing no appreciable differences were excluded ($p \geq 0.1$), yielding 44 features derived from 9 base parameters carried forward for recovery metric computation.

Recovery was quantified using a composite kinematic score (CKS), which performs principal component analysis on extracted gait features and calculates Euclidean distance by projecting data onto the line connecting baseline and acute injury (7 DPI) centroids. CKS scores were normalized to a 0–100 scale, where a score of 0 corresponds to acutely injured animals (vehicle, 7 DPI) and 100 to healthy baseline animals (0 DPI):

$$CKS = 100 \times \frac{score_i - \mu_{injured}}{\mu_{baseline} - \mu_{injured}}$$

To verify the method independence of the results, three additional recovery metrics (Mahalanobis Distance, kNN Baseline Distance, and Ridge Regression Score) were developed. The Mahalanobis Distance metric was computed as the distance from each sample to the healthy baseline distribution in the 10-dimensional PC space, providing a covariance-adjusted multivariate measure of deviation from healthy locomotion. A non-parametric recovery metric, the k-Nearest Neighbor Baseline Distance (kNN), was computed from Euclidean distances in the full 44-feature space. For each post-injury sample, distance to its 6 nearest baseline animal neighbors ($k \approx \sqrt{N}$, $N = 38$ baseline animals) was averaged and converted to a similarity score as $1/(1 + \bar{d})$, where $\bar{d}$ denotes the mean distance to the k nearest neighbors. A supervised Ridge Regression Score was derived from a Ridge regression model trained to distinguish healthy baseline animals (target = 1.0) from acutely injured vehicle animals at 7 DPI (target = 0.0). The model incorporated the 44 selected kinematic features, their interactions with two binary treatment indicator variables, and the indicator variables themselves. The regularization parameter was selected by maximizing mean $R^2$ across stratified 5-fold cross-validation ($\alpha \in \{1, 10, 50, 100, 200, 500, 1000, 2000\}$). All three metrics were normalized to a common 0–100 scale anchored at the mean score of acutely injured animals (vehicle, 7 DPI; score = 0) and the mean score of healthy baseline animals (0 DPI; score = 100): $\text{score} = 100 \times (\text{score}_i - \mu\text{injured}) / (\mu\text{baseline} - \mu\text{injured})$.

**Kinematic trace analysis**

Kinematic trace analyses were performed in MATLAB (R2025b). For each video frame, the XY position of the right forelimb toe was expressed relative to the right ear by subtracting the ear coordinates, yielding a two-dimensional displacement vector that captures limb trajectory independently of whole-body forward progression. The resulting time series were smoothed with a five-frame moving average. Gait cycles were segmented by detecting local maxima in the X-displacement signal with a minimum inter-peak separation of 20 frames. Each cycle was time-normalized to 100 equally spaced points by PCHIP interpolation and centered prior to averaging. Trial-level traces were averaged across trials within each animal (5 trials/animal/session) and used to calculate group-level mean toe-to-ear traces ± SEM. Finally, group traces were circularly normalized and aligned to facilitate visual comparisons of weight-bearing behavior between groups.

**Spinal cord histology and immunostaining**

At 42 DPI, rodents were deeply anesthetized using 3% isoflurane until animals showed no response to physical stimuli and were transcardially perfused with ice-cold PBS (150–250 mL), followed by ice-cold 4% PFA (200–300 mL). The spinal cord was dissected and post-fixed overnight in 4% PFA at 4°C, then transferred to PBS and kept at 4°C prior to sectioning. The injury epicenter was identified under a dissecting microscope by its characteristic brownish discoloration, and a 1 cm segment centered on the epicenter (0.5 cm rostral and 0.5 cm caudal) was isolated. The tissue segment was embedded in 2% agarose/PBS and sectioned at 100 μm in the transverse plane using a Compresstome vibrating microtome (VF-510-0Z; Precisionary Instruments, Natick, MA, USA). Serial sections were collected into a 24-well plate containing PBS (five sections per well), with rostral-to-caudal order, and maintained at 4°C until immunostaining.

Tissue sections were washed in PBS and transferred to 2 mL microcentrifuge tubes containing 750 μL of blocking buffer (2.5% bovine serum albumin and 0.1% Triton X-100 in PBS). Tissues were blocked for 1 h at room temperature with gentle rotation. The sections were treated with glial fibrillary acidic protein primary antibody (GFAP; Dako, Z0334, RRID:AB_10013382) diluted 1:1,000 in blocking buffer overnight at 4°C with gentle rotation. Tissues were transferred to a 12-well plate fitted with mesh inserts and washed 3 × 10 min in PBS. Sections were incubated with goat anti-rabbit IgG conjugated to Alexa Fluor 488 (Thermo Fisher Scientific, A11034, RRID:AB_2576217; 1:1,000) in blocking buffer for 2 h at room temperature with gentle rotation. Nuclear counterstaining was performed with Hoechst 33342 (Invitrogen, H3570) at 1:1,000 for 5 min, followed by 3 × 10 min PBS washes. Sections were mounted onto glass microscope slides (VWR Micro Slides, 48311-703), coverslipped with ProLong Gold Antifade Mountant (Invitrogen, P36931), and allowed to cure in the dark for a minimum of 6 h before imaging. Fluorescence images were acquired using an Andor Dragonfly spinning disk confocal microscope equipped with a 10X objective.

**Quantification of lesion size**

Histological images were analyzed using a custom, deep learning–assisted image analysis pipeline implemented in Python using the PyTorch framework. A U-Net model was trained to segment the spinal cord boundary from the background, and model-assisted segmentation was used to classify tissue regions as injured, uninjured, or lesioned. Models were trained using manually annotated regions of interest generated with a custom annotation tool, with training performed on GPUs.

For image analysis, model predictions were generated using tiled inference with 1,024-pixel patches and 25% overlap. The initial outputs were subsequently refined using custom image-processing steps to improve tissue boundary detection, repair discontinuities in damaged spinal cord sections, partition the cord into injured and uninjured regions, and identify lesion cavities as regions of missing or low-intensity tissue within the injured hemicord. Segmentation overlays showing the spinal cord boundary, injured–uninjured partition, and lesion mask were visually inspected as part of the quality control process before quantitative analysis.

Lesion size (%) was calculated for each section as the lesion area divided by the total spinal cord cross-sectional area. Values were averaged across sections for each animal, and treatment groups were summarized as median values with 95% confidence intervals using GraphPad Prism.

**Assessment of astrocyte and microglial reactivity**

Adult male Sprague-Dawley rats (6-7 weeks old) were randomly assigned to two experimental groups (n = 7 per group): vehicle control and TMP-316 at ~1 mg/kg. IT catheterization and drug delivery were performed at the L6–S1 interlaminar space as previously described by Elwardany et al.[14] Following surgery, animals were single housed, placed on a heating pad for recovery, and administered a single dose of Buprenorphine-SR (Ethiqa XR; 0.65 mg/kg, subcutaneous) for post-operative analgesia. At 14 days post-treatment (sufficient for astrogliosis and microglial activation to manifest if induced), rats were deeply anesthetized with ketamine (225 mg/kg; Dechra) and xylazine (30 mg/kg; Rompun, Dechra) and transcardially perfused with ice-cold PBS, followed by ice-cold 4% PFA. The lumbar spinal cord was dissected and post-fixed overnight in 4% PFA at 4°C, followed by cryoprotection in 30% sucrose/PBS for 72 h. Samples were frozen in Epredia™ M-1 Embedding Matrix (Fisher Scientific) using a dry ice slurry. Tissue was cryosectioned at 16 µm (Leica CM3050S) and mounted onto glass microscope slides (VWR Micro Slides, 48311-703) before staining.

Slides containing rostral (L1–L2) and caudal (L4–L5) lumbar spinal cord sections were equilibrated at room temperature for 1 h, followed by 3 × 5 min PBS washes. Sections were blocked in 5% normal goat serum in 0.3% Triton X-100/PBS (PBS-T) for 45 min at room temperature. Sections were incubated overnight at 4°C with primary antibodies against Iba-1 (Abcam, ab178846, RRID:AB_2636859; 1:500) and GFAP (Abcam, ab4674, RRID:AB_304558; 1:1,000). Sections were then incubated with goat anti-chicken Alexa Fluor 568 (Abcam; 1:500) and goat anti-rabbit Alexa Fluor 647 (Abcam; 1:500) for 1 h at room temperature, washed 3 × 5 min in PBS-T, then coverslipped with ProLong Gold Antifade Mountant (Invitrogen, P36931), and allowed to cure in the dark overnight. Slides were imaged on a Nikon Eclipse Ti-2 microscope.

GFAP fluorescence intensity and Iba-1-positive (Iba-1+) cell counts were quantified in ImageJ (v2.16). Briefly, the spinal cord section was delineated as the region of interest (ROI), and its area measured. Images were converted to 8-bit grayscale and thresholded using a common threshold value across all images. The mean fluorescence intensity within the ROI was then measured to quantify GFAP immunoreactivity. The number of Iba-1+ cells was quantified using ImageJ's Analyze Particles function, then normalized to ROI area to yield Iba-1+ cell density. Values were averaged across sections for each animal, with a minimum of two sections per animal.

### References


1. Awada, B. *et al.* Phenotypic Screening with Primary and Human iPSC-derived Neurons. *Assay Guidance Manual* https://www.ncbi.nlm.nih.gov/books/NBK619698/ (2025).
2. Schmittgen, T. D. & Livak, K. J. Analyzing real-time PCR data by the comparative C(T) method. *Nat. Protoc.* **3**, 1101–1108 (2008).
3. Nagendran, T., Poole, V., Harris, J. & Taylor, A. M. Use of Pre-Assembled Plastic Microfluidic Chips for Compartmentalizing Primary Murine Neurons. *J. Vis. Exp.* **2018**, (2018).

4. Wu, C., Schulte, J., Sepp, K. J., Littleton, J. T. & Hong, P. Automatic robust neurite detection and morphological analysis of neuronal cell cultures in high-content screening. *Neuroinformatics* **8**, 83–100 (2010).

5. BLISS, C. I. THE TOXICITY OF POISONS APPLIED JOINTLY. *Annals of Applied Biology* **26**, 585–615 (1939).

6. Manning, G., Whyte, D. B., Martinez, R., Hunter, T. & Sudarsanam, S. The protein kinase complement of the human genome. *Science* **298**, 1912–34 (2002).

7. Kumar, S. *et al.* TimeTree 5: An Expanded Resource for Species Divergence Times. *Mol. Biol. Evol.* **39**, (2022).

8. Passaro, S. *et al.* Boltz-2: Towards Accurate and Efficient Binding Affinity Prediction. *bioRxiv* https://doi.org/10.1101/2025.06.14.659707 (2025) doi:10.1101/2025.06.14.659707.

9. Tauriello, G. *et al.* ModelArchive: A Deposition Database for Computational Macromolecular Structural Models. *J. Mol. Biol.* **437**, 168996 (2025).

10. Anastassiadis, T., Deacon, S. W., Devarajan, K., Ma, H. & Peterson, J. R. Comprehensive assay of kinase catalytic activity reveals features of kinase inhibitor selectivity. *Nat. Biotechnol.* **29**, 1039–45 (2011).

11. Eid, S., Turk, S., Volkamer, A., Rippmann, F. & Fulle, S. KinMap: a web-based tool for interactive navigation through human kinome data. *BMC Bioinformatics 2017 18:1* **18**, 16- (2017).

12. Perez-Riverol, Y. *et al.* The PRIDE database at 20 years: 2025 update. *Nucleic Acids Res.* **53**, D543–D553 (2025).

13. Christ, M., Braun, N., Neuffer, J. & Kempa-Liehr, A. W. Time Series FeatuRe Extraction on basis of Scalable Hypothesis tests (tsfresh – A Python package). *Neurocomputing* **307**, 72–77 (2018).

14. Elwardany, O. S. *et al.* Lumbar intrathecal catheterization in rats targeting the cerebral cortex: a drug delivery method and validation. *bioRxiv* 2026.06.01.727192 (2026) doi:10.64898/2026.06.01.727192.

Supplementary Figure 1

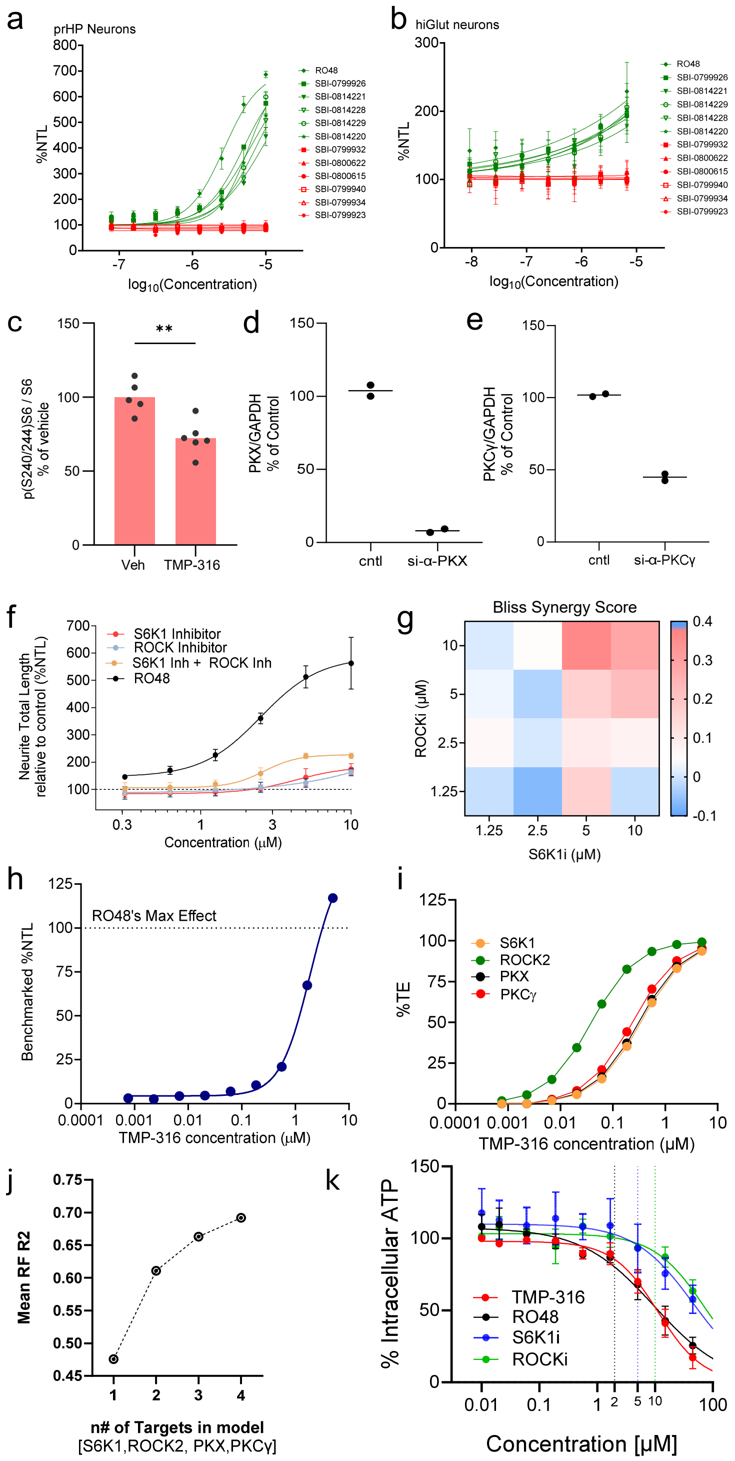

**Supplementary Figure 1 | Extended in vitro analyses of program compounds**

**(a)** Neurite outgrowth (%NTL) of prHP neurons (n = 2–5 technical replicates) and (b) hiGLUT neurons (n = 6 replicates) treated with active and inactive RO48 analogs. Data are presented as mean ± SD. **(c)** p(S240/S244)S6/S6 levels quantified by Western blot from cortical lysates of rats treated with vehicle or TMP-316 (1 mg/kg). Data are presented as mean showing individual data points (n = 5–6 animals per group; unpaired t-test, ** $p < 0.01$). mRNA expression of PKX **(d)** and PKCγ **(e)** in prHP neurons treated with scramble (cntl) or targeting siRNA (α-PKX or α-PKCγ, respectively). Data are presented as mean ± SD (n= 2 biological replicates). **(f)** Neurite outgrowth (%NTL) of prHP neurons treated with S6K1i, ROCKi, S6K1i+ROCKi, or RO48 across concentrations ranging from 0.3 – 10 μM. Data are presented as mean ± SD (n = 3 technical replicates). **(g)** Bliss synergy scores calculated for S6K1i vs ROCKi treatment across a range of concentrations (1.25, 2.5, 5, 10 μM). Higher values indicate synergistic enhancement of neurite outgrowth. **(h)** TMP-316's benchmarked %NTL in prHP neurons treated with RO48. **(i)** TMP-316's target engagement (%TE) curves with S6K1, ROCK2, PKX, and PKCγ. **(j)** Random forest model performance (mean RF $R^2$) as a function of the number of targets included (S6K1, ROCK2, PKX, PKCγ). Each point represents an independent model fit. **(k)** CellTiter-Glo® Cell Viability Assay dose-response of E18 prHP neurons incubated for 48 hours with TMP-316, RO48, ROCKi, or S6K1i normalized to DMSO control. Shown are mean ± SD, N = 2-3 biological replicates with 2 technical replicates in each.

Supplementary Figure 2

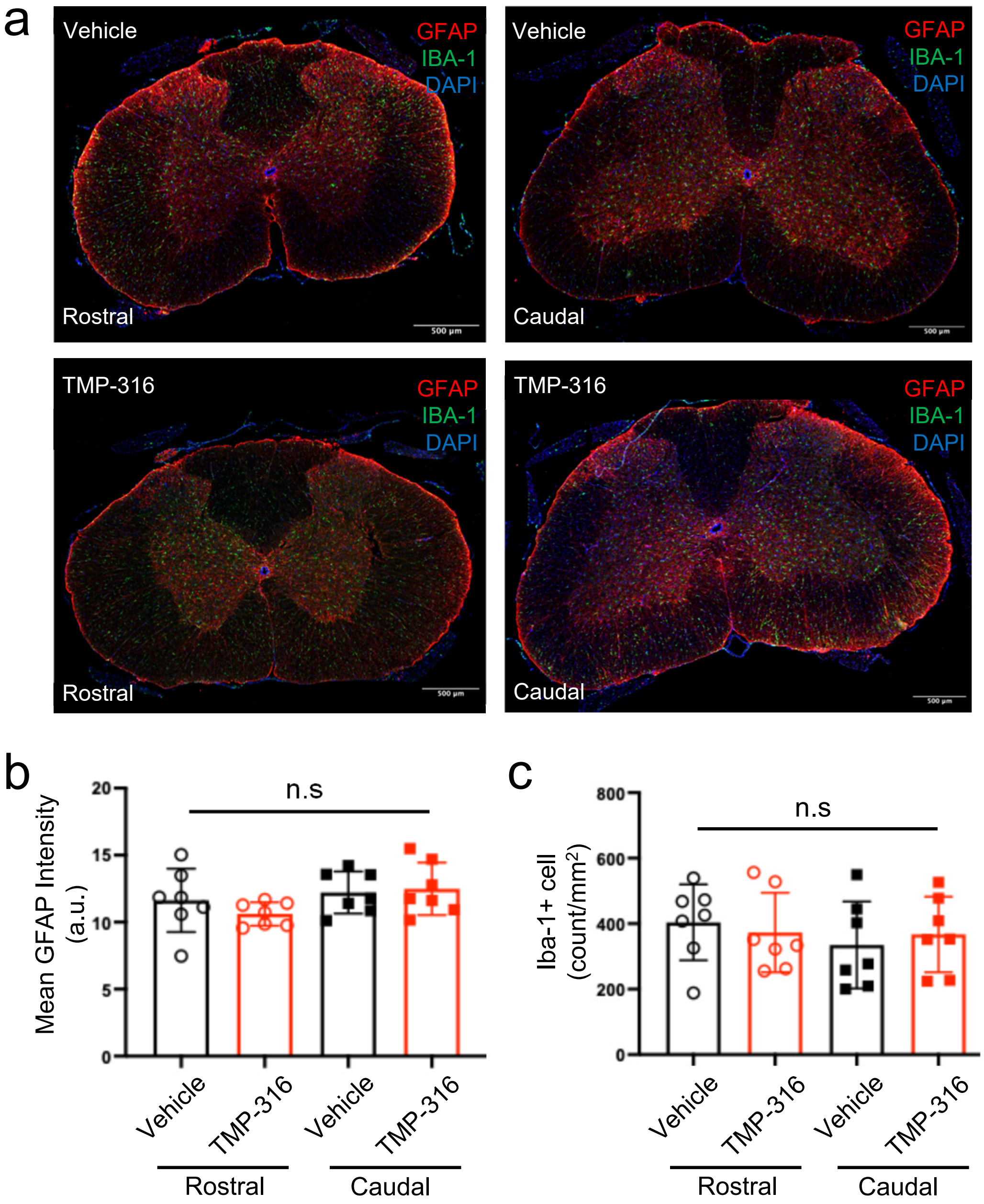

**Supplementary Figure 2 | Astrocyte and microglial markers are unchanged after intrathecal delivery of TMP-316**

Transverse sections of the rostral (L1–L2) and caudal (L4–L5) lumbar spinal cord collected 14 days after intrathecal (IT) delivery of vehicle or TMP-316 (~1 mg/kg). (**a**) Representative fluorescence images of sections immunostained for GFAP (red), Iba-1 (green), and DAPI (blue). Scale bars: 500 µm. (**b**) Quantification of mean GFAP fluorescence intensity. (**c**) Quantification of Iba-1+ cell density. Data are presented as mean ± SEM (N = 7 per group; one-way ANOVA with Tukey's multiple comparisons test).

Supplementary Figure 3

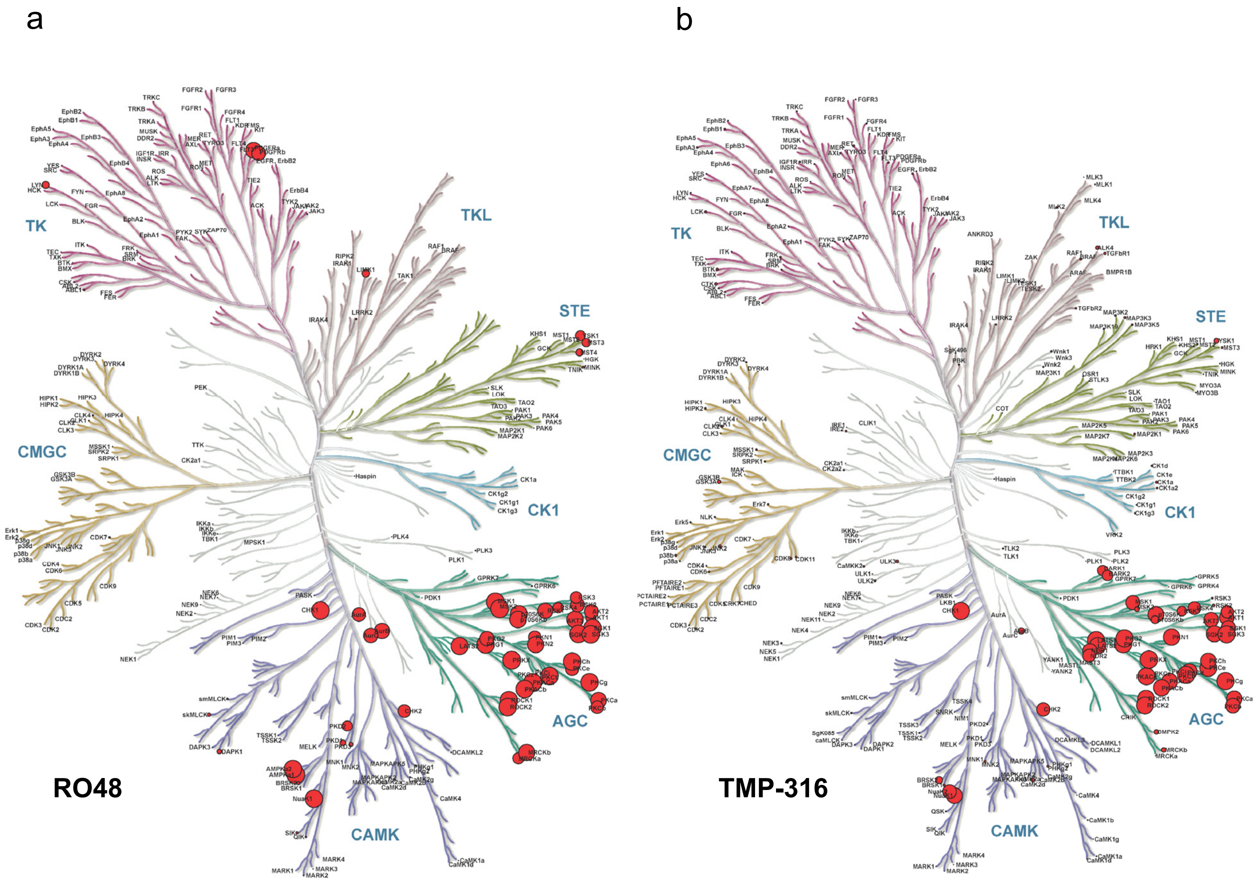

**Supplementary Figure 3 | Broad kinome profiling of RO48 and TMP-316**

Node size is scaled within each compound to its maximum observed inhibition, so distribution across the kinome is comparable. Only kinases with assay data are annotated on each map. (a) RO48, profiled at 100 nM at Km ATP (maximum inhibition ~100%). (b) TMP-316, profiled at 500 nM in 1 mM ATP (maximum inhibition ~100%). Several non-AGC kinases inhibited by RO48 are not engaged by TMP-316. Illustration courtesy of Cell Signaling Technology, Inc. (www.cellsignal.com).

Supplementary Figure 4

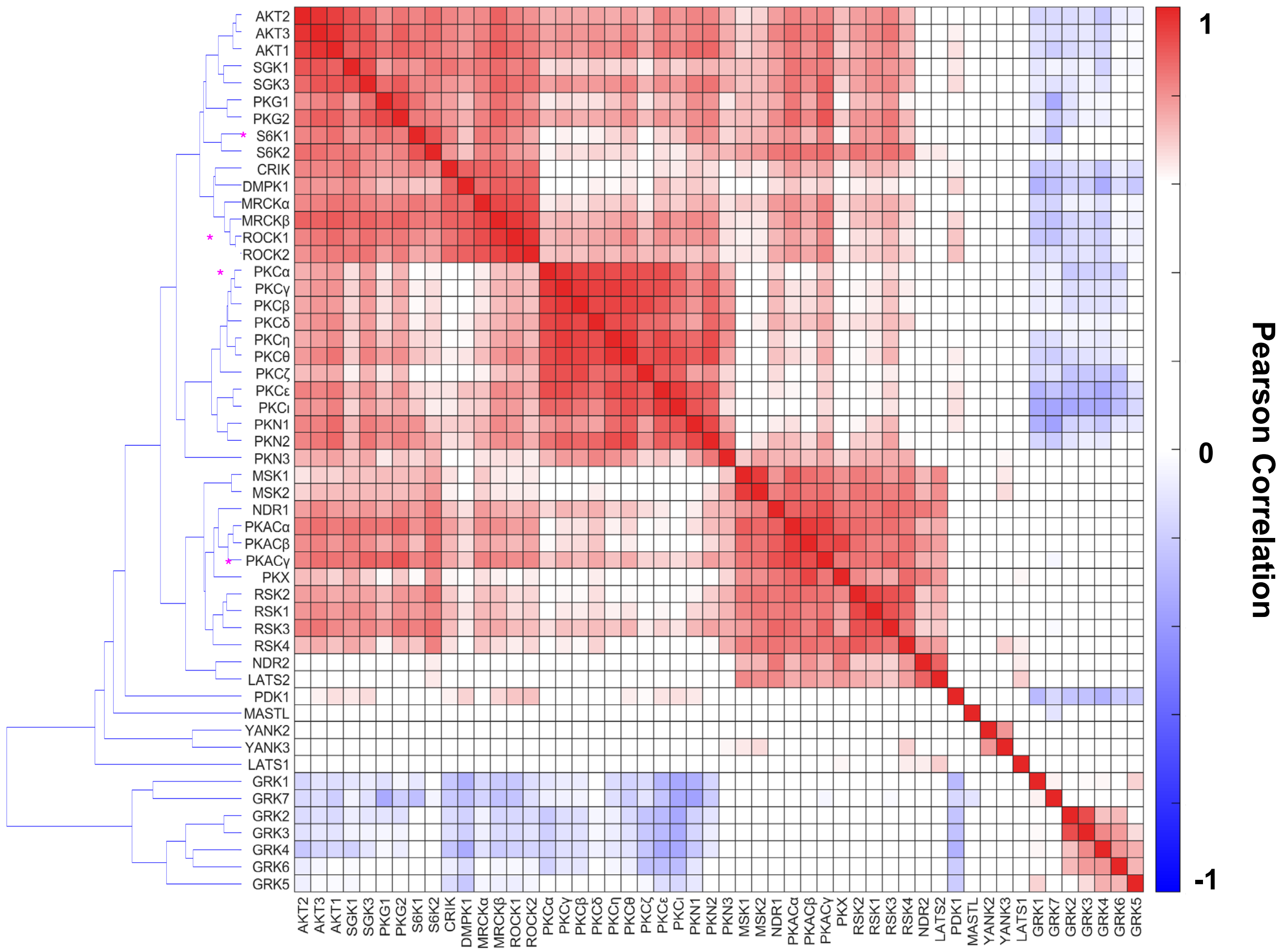

**Supplementary Figure 4 | Full Annotated Substrate Sequence Preference Profiles**

Shown is the pairwise Pearson correlation of substrate sequence preference profiles across AGC kinases from Figure 3b, with full kinase annotations.

Supplementary Figure 5

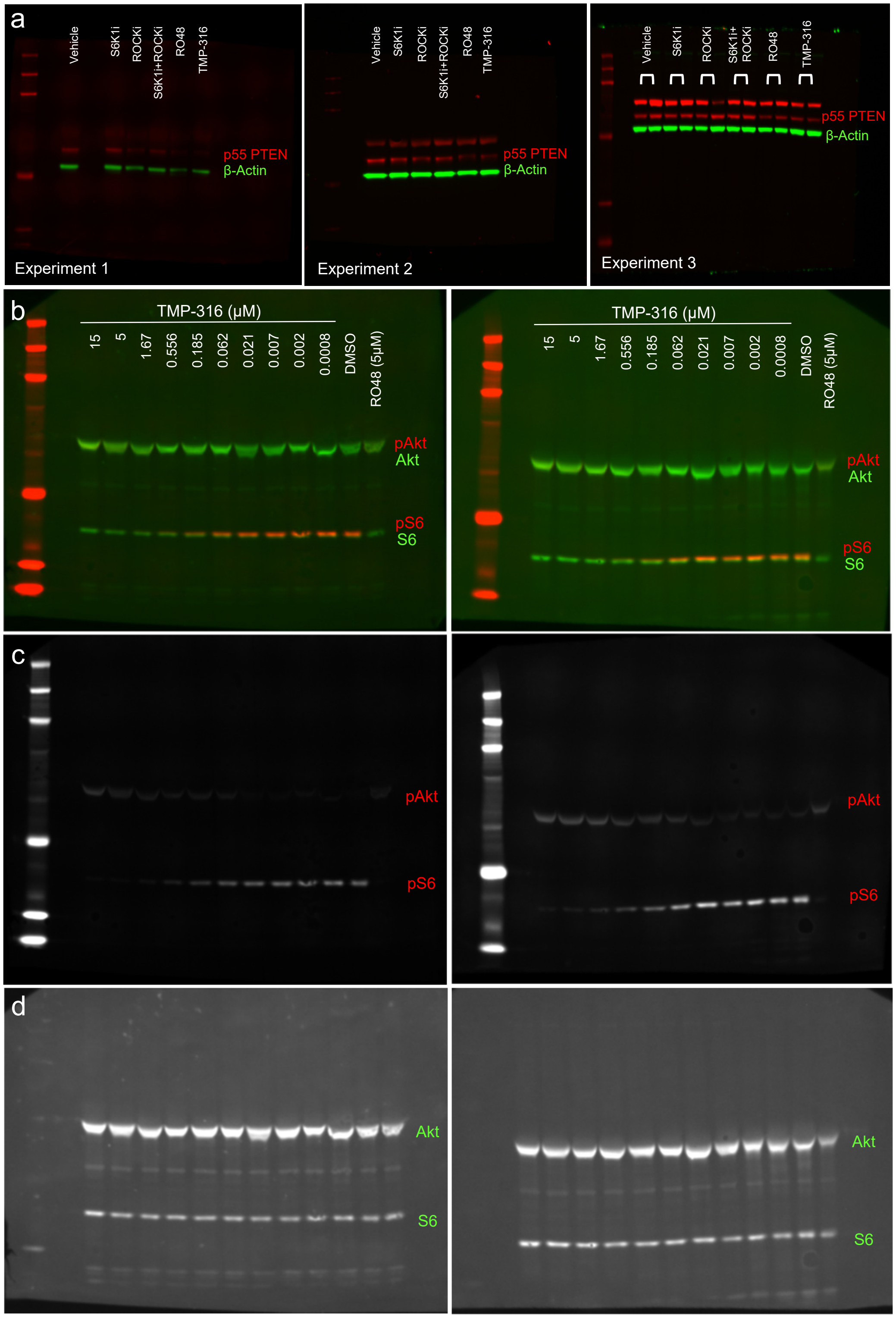

**Supplementary Figure 5 | Western Blot analysis of PI3K/PTEN signaling markers**

**(a)** Immunoblots from three independent experiments showing levels of p55 PTEN (red) and the housekeeping protein β-actin (green) in prHP neurons treated with vehicle, S6K1i (5 μM), ROCKi (10 μM), S6K1i + ROCKi, RO48 (2 μM), or TMP-316 (2 μM). **(b)** Immunoblots from two technical replicates showing levels of p(S473)Akt, total Akt, p(S240/244)S6, and total S6 in prHP neurons treated with a range of TMP-316 concentrations (0.8 nM–15 μM), alongside vehicle and RO48 (5 μM) controls. **(c–d)** Grayscale extractions of the red (phospho) and green (total protein) channels corresponding to panel (b), shown for each replicate.

Supplementary Figure 6

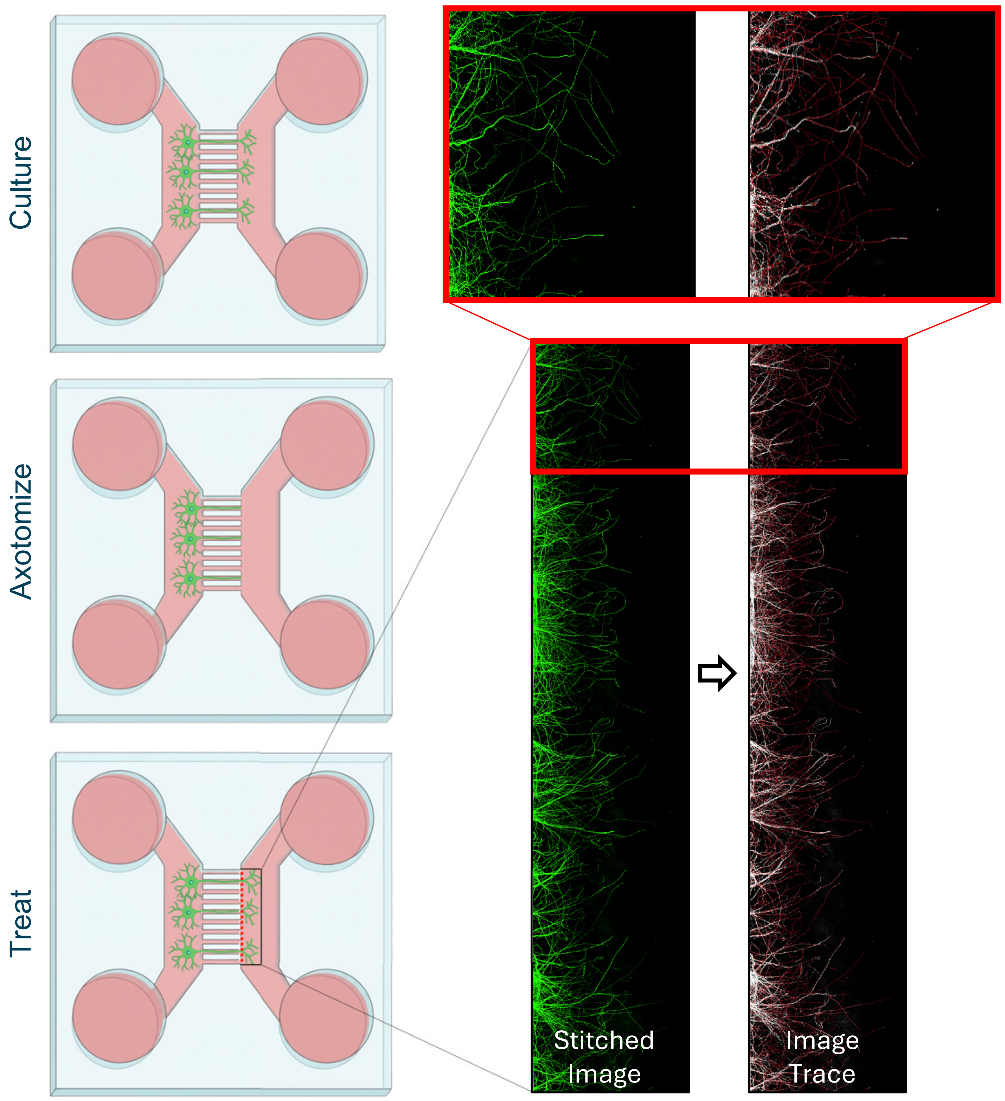

**Supplementary Figure 6 | Microfluidic axotomy assay workflow and image analysis**

Neurons were cultured in microfluidic chambers for 8 DIV, axotomized by vacuum-mediated severing, and treated for 48 h. Chambers were then stained for βIII-tubulin (green) to visualize axons and imaged at 1 μm z-intervals. Image stacks were stitched to generate maximum-intensity projections, which were then processed to produce red tracing overlays used to quantify axon regrowth.

Supplementary Figure 7

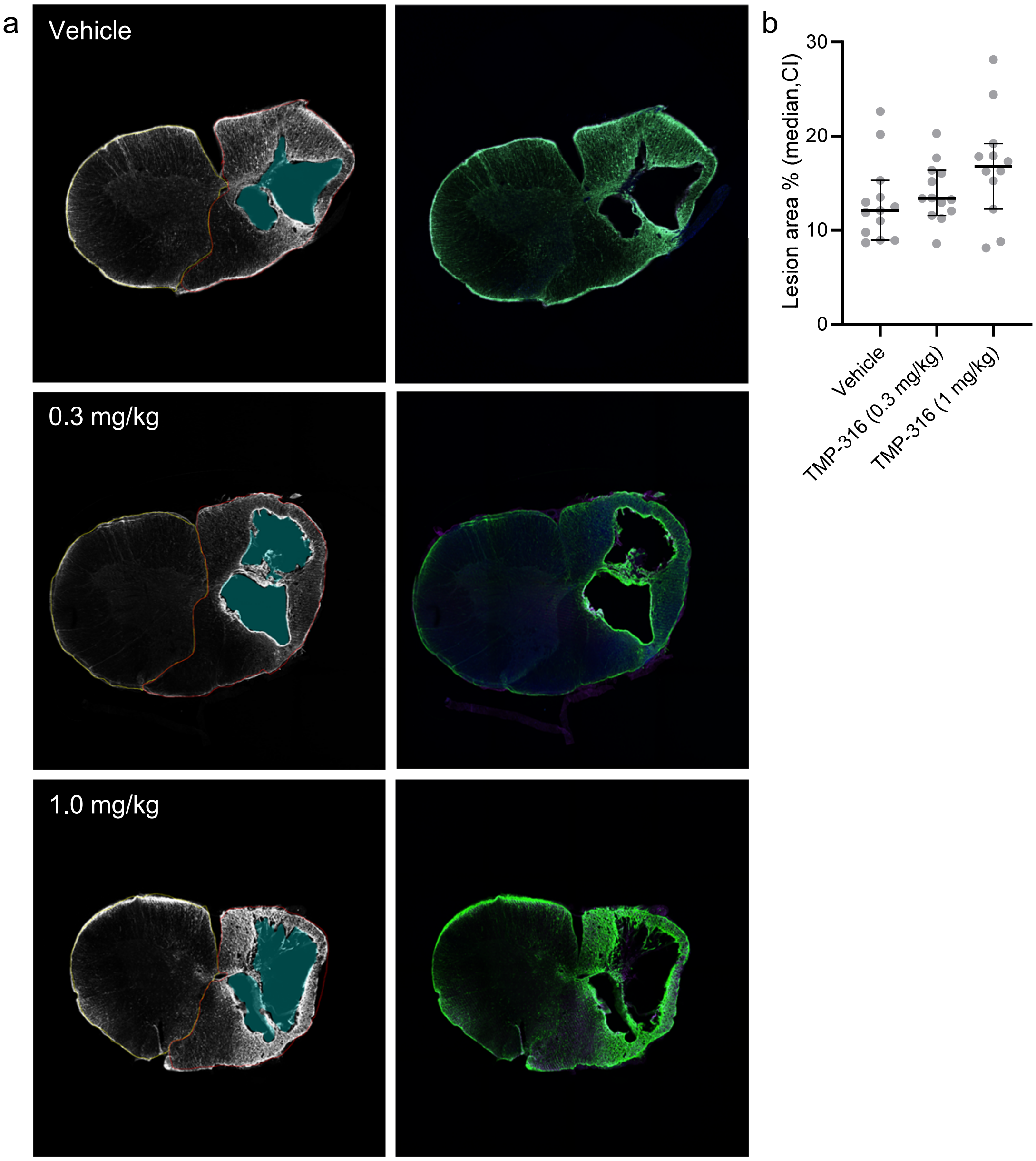

**Supplementary Figure 7 | Spinal cord lesion area analysis**

**(a)** Transverse spinal cord sections taken from the C5–C6 injury epicenter at 42 DPI from rats treated with vehicle, TMP-316 0.3 mg/kg, or TMP-316 1 mg/kg. Right panels show representative fluorescence images of sections immunostained for GFAP (green) with Hoechst counterstaining (blue). Left panels show corresponding U-Net segmentation masks classifying tissue as uninjured, injured, or lesion, with lesion regions outlined and filled according to an automated pipeline where blue indicates lesion area, red denotes injured spinal cord, and yellow designates uninjured spinal cord. **(b)** Lesion area (%) quantified per animal as lesion area divided by total spinal cord cross-sectional area, averaged across all sections spanning the injury epicenter. Data are presented as median values with 95% confidence intervals (N = 12–13 animals per group).

Supplementary Figure 8

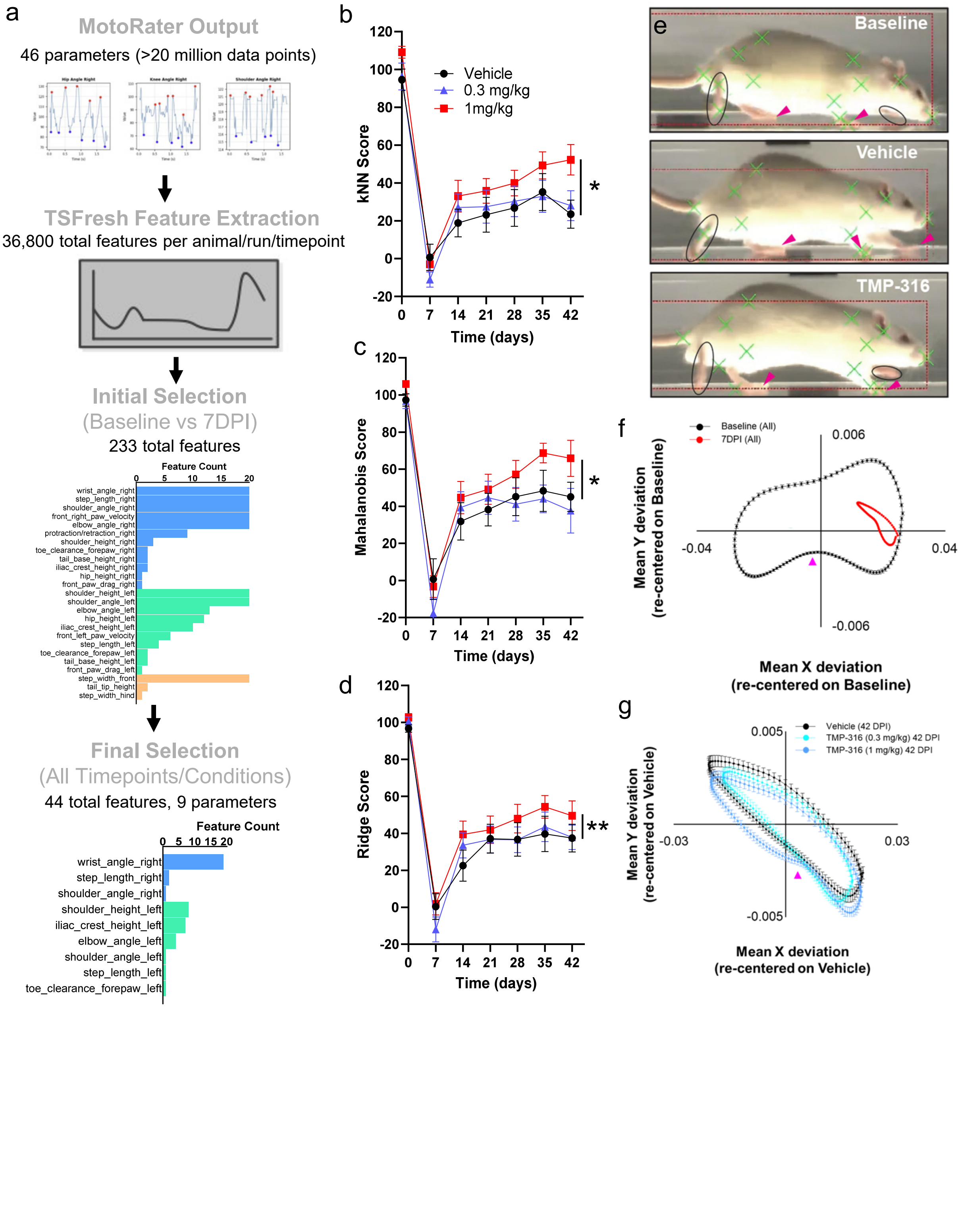

**Supplementary Figure 8 | Feature-driven gait analysis and model scoring highlight motor-function rescue following TMP-316 treatment**

**(a)** Workflow for processing MotoRater time-series data. Over 20 million data points from 46 base parameters were subjected to TSFresh feature extraction, yielding ~36,800 features per animal per run. Initial filtering (baseline vs 7 DPI) identified 233 features across multiple gait parameters, followed by treatment-responsive filtering across all timepoints, resulting in 44 final features derived from 9 base parameters. **(b–d)** Recovery trajectories computed using three independent metrics: k-Nearest Neighbor (kNN) Score **(b)**, Mahalanobis Score **(c)**, and Ridge Regression Score **(d)**. Scores were normalized to a 0–100 scale anchored at baseline (0 DPI) and acutely injured vehicle animals (7 DPI). Data are presented as mean ± SEM for vehicle, TMP-316 0.3 mg/kg, and TMP-316 1 mg/kg groups (N = 12–13 per group; One-Way Repeated Measures ANOVA with Dunnett's multiple comparisons test; * $p < 0.05$, ** $p < 0.01$). **(e)** Representative MotoRater frames with joint-tracking overlays (green) at baseline, 42 DPI vehicle, and 42 DPI TMP-316 1 mg/kg. Pink arrowheads indicate foot–ground contact. **(f)** Consensus toe-to-ear deviation traces of the right forelimb, comparing control animals (baseline) and acutely injured animals at 7 DPI. Traces were averaged across trials and animals, and plotted as group mean ± SEM. **(g)** Consensus toe-to-ear deviation traces, comparing vehicle, TMP-316 0.3 mg/kg, and TMP-316 1 mg/kg groups. Traces were averaged across trials and animals and plotted as group mean ± SEM (N = 12–13 animals per group). Pink arrowhead indicates point of maximum weight-bearing, identified by minimum toe-to-ear distance during stance phase.

Supplementary Figure 9

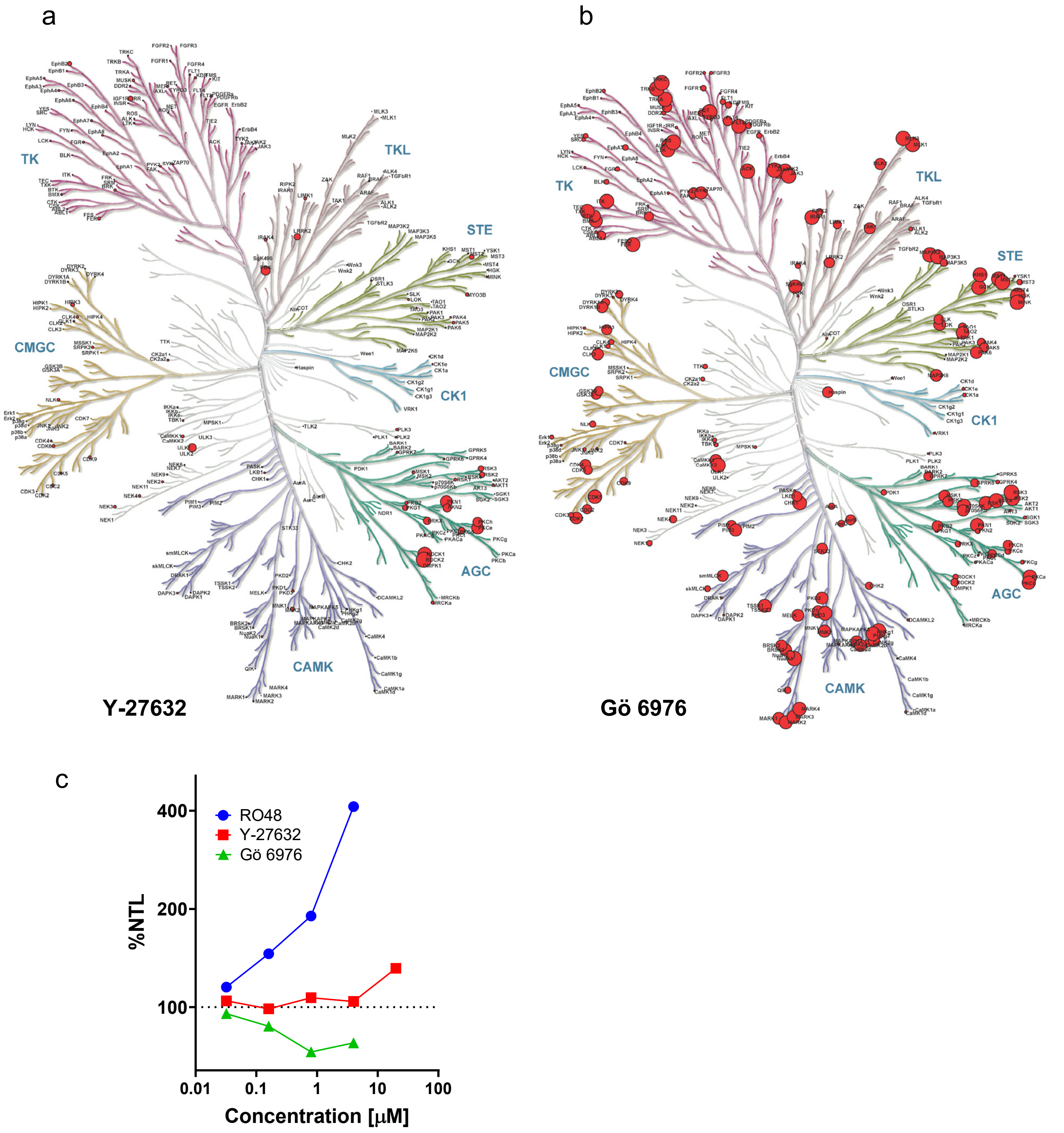

**Supplementary Figure 9 | Broad kinome profiling and neurite outgrowth (prHP) activity of Y-27632 and Gö 6976**

Node size is scaled within each compound to its maximum observed inhibition, so distribution across the kinome is comparable between maps but absolute node sizes are not. Only kinases with assay data are annotated on each map. (a, b) Y-27632 and Gö 6976, profiled at 500 nM in 10 μM ATP; data from Anastassiadis et al. (2011). (c) Effect of the indicated compounds on neurite outgrowth in the prHP assay (data adapted from Al-Ali et al., 2015).
